# Evolutionary analysis of conserved non-coding elements subsequent to whole-genome duplication in opium poppy

**DOI:** 10.1101/2023.05.06.539671

**Authors:** Yu Xu, Stephen J. Bush, Xinyi Yang, Linfeng Xu, Bo Wang, Kai Ye

**Affiliations:** School of Life Science and Technology, Xi’an Jiaotong University, Xi’an, Shaanxi, China; School of Automation Science and Engineering, Faculty of Electronic and Information Engineering, Xi’an Jiaotong University, Xi’an, Shaanxi, China; MOE Key Lab for Intelligent Networks & Networks Security, Faculty of Electronic and Information Engineering, Xi’an Jiaotong University, Xi’an, Shaanxi, China; Genome Institute, the First Affiliated Hospital of Xi’an Jiaotong University, Xi’an, Shaanxi, China

**Keywords:** Conserved non-coding elements, transcriptomic evolution, whole-genome duplication, fractionation, BIA gene cluster

## Abstract

Whole-genome duplication (WGD) leads to the duplication of both coding and non-coding sequences within an organism’s genome, providing an abundant supply of genetic material that can drive evolution, ultimately contributing to plant adaptation and speciation. Although non-coding sequences contain numerous regulatory elements, they have been understudied compared to coding sequences. In order to address this gap, we explored the evolutionary patterns of regulatory sequences, coding sequences, and transcriptomes using conserved non-coding elements (CNEs) as regulatory element proxies following the recent WGD event in opium poppy (*Papaver somniferum*). Our results showed similar evolutionary patterns in subgenomes of regulatory and coding sequences. Specifically, the biased or unbiased retention of coding sequences reflected the same pattern as retention levels in regulatory sequences. Further, the divergence of gene expression patterns mediated by regulatory element variations occurred at a more rapid pace than that of gene coding sequences. However, gene losses were purportedly dependent on relaxed selection pressure in coding sequences. Specifically, the rapid evolution of tissue-specific benzylisoquinoline alkaloids production in *P. somniferum* was associated with regulatory element changes. The origin of a novel stem-specific ACR, which utilized ancestral cis-elements as templates, is likely to be linked to the evolutionary trajectory behind the transition of the *PSMT1-CYP719A21* cluster from high levels of expression solely in *P. rhoeas* root tissue to its elevated expression in *P. somniferum* stem tissue. Our findings demonstrate that rapid regulatory element evolution can contribute to the emergence of new phenotypes and provide valuable insights into the high evolvability of regulatory elements.

**Significance Statement:** This study demonstrates that rapid evolution of regulatory elements can drive the emergence of novel phenotypes in plants. Our investigation, in particular, revealed that the evolution of stem-specific high expression patterns of BIAs genes in *P. somniferum* was linked to rapid changes in regulatory elements.

## Introduction

Understanding the molecular mechanisms of phenotypic adaptation and speciation is a fundamental objective in the field of biology (Romero *et al*., 2012). Whole-genome duplication (WGD) events are commonly occurring in the evolution of plants, leading to the doubling of both coding and non-coding sequences (Liang and Schnable, 2018, Qiao *et al*., 2019). Compared to variations in coding regions, variations in non-coding regions with abundant regulatory elements can alter gene expression patterns without affecting protein structure and function and are emphasized as more common and important for evolution (King and Wilson, 1975, Wray, 2007, Carroll, 2008, Necsulea and Kaessmann, 2014). However, there is a lack of systematic and comprehensive genome-wide studies elucidating the role of non-coding DNA variations in enhancing transcriptomic and phenotypic evolution in plants (Necsulea and Kaessmann, 2014, Yocca and Edger, 2022).

The detection of useful non-coding regulatory elements still poses a significant challenge in plant science, particularly in cases where comprehensive ENCODE-scale data annotation is lacking (Song *et al*., 2021). To tackle this challenge, comparative genomics approaches based on evolutionary principles have been adopted to identify conserved non-coding elements (CNEs). These elements, usually at least 15 bp long and subjected to strong purifying selection and sequence constraint, demonstrate a high degree of conservation across species, as reported by Turco et al. (Turco *et al*., 2013) and Yocca et al. (Yocca *et al*., 2021). CNEs correspond with functional genomics signatures, such as transcription factor binding sites (TFBSs) and accessible chromatin regions (ACRs), and their variations are linked with alterations in gene expression patterns (Turco *et al*., 2013, Song *et al*., 2021, Yocca *et al*., 2021, Yocca and Edger, 2022). Consequently, CNEs serve as an effective surrogate for cis-regulatory elements (CREs) to uncover novel mechanisms of transcriptomic and phenotypic evolution.

Recurrent duplication events and dense repeats pose a significant challenge in plant science, as they hinder the availability of high-quality plant genomes can be used for comparative analysis. However, recent publications of high-quality *Papaver* genomes and transcriptomes have provided valuable data to facilitate systematic study and comparison research (Yang *et al*., 2021, Xu *et al*., 2022). The opium poppy (*Papaver somniferum*) is one of the oldest and most important medicinal plants worldwide, producing several pharmaceutically important benzylisoquinoline alkaloids (BIAs), including noscapine, morphine and codeine (Beaudoin and Facchini, 2014). *P. somniferum* and *P. rhoeas* (common poppy) diverged around 7.7 million years ago (Mya), after which *P. somniferum* underwent a lineage-specific WGD, leading to the formation of the stem-specificity BIA gene cluster (Yang *et al*., 2021). The innovation in BIA gene cluster formation is accompanied by rapid alterations in the expression profiles of BIAs biosynthetic genes and increased production of both noscapine and morphine. This spectacular adaptation and evolutionary trajectory position the *Papaver* species as an excellent model system to investigate a variety of pertinent evolutionary inquiries.

After WGD, gene loss or fractionation occurs, which was initially studied in terms of redundant gene loss in newly duplicated genomic regions, is now extends to non-coding or even epigenetic fields (Schnable James *et al*., 2011, Edger *et al*., 2017, Zhao *et al*., 2017, Liang and Schnable, 2018, Miao *et al*., 2020, Zhang *et al*., 2021). As *P. rhoeas* is closely related to *P. somniferum* but did not undergo the *P. somniferum*-specific WGD event, it can be considered as a pre-WGD ancestor of *P. somniferum* and used to distinguish the two subgenomes after the WGD event. Therefore, by using genes as anchors and assuming the *P. rhoeas* genome organisation as pre-WGD ancestor status, a systematical comparison of non-coding sequence variations between subgenomes can be conducted.

In this study, we reconstructed two subgenomes of *P. somniferum* and identified CNEs based on the syntenic relationship between these subgenomes and the *P. rhoeas* genome. These data, combined with transcriptomic and epigenomic landscapes (previously constructed in our laboratory for these non-model medicinal crops across multiple tissues (Yang *et al*., 2021, Jia *et al*., 2023)), were used to achieve the following four objectives: (1) to investigate the evolutionary trajectory and relationship between coding sequences and non-coding sequences in two subgenomes of *P. somniferum*; (2) to elucidate the impact of variations (presence or absence) conserved non-coding elements on the evolution of gene expression patterns; (3) to compare the divergent rates of evolution between gene products and gene expression patterns; (4) to shed light on the evolution of transcriptional profiles of BIA gene cluster through analysis of conserved non-coding elements.

## Results

### Unbiased fractionation of the two subgenomes of opium poppy

The genome of *P. rhoeas* could be treated as two diploid subgenomes present in the tetraploid ancestor of *P. somniferum* (Yang *et al*., 2021). A total of 11,765 WGD gene pairs and 21,848 singletons (genes with no duplicate) were identified after removing tandemly duplicated genes in *P. somniferum*. These genes were used to identify syntenic relationships between *P. somniferum* and its outgroup diploid relative *P. rhoeas* (Yang *et al*., 2021). The genes with syntenic homologous genes are known as “syntelogs” based on the classical definition (Zhao *et al*., 2017). A majority (68.5%, or 8,064) of the duplicated gene pairs had syntelogs in the *P. rhoeas* genome, while only a minority (19.4%, or 4,235) of the singleton genes had syntelogs in the *P. rhoeas* genome. Assuming that the genes with syntelogs represent accurately annotated homologs, the ratio of singletons to duplicated gene pairs in *P. somniferum* was estimated to be 1:1.9. By contrast, the biased fractionated genome of maize had a ratio of 1:0.39, while the unbiased fractionated genome of soybean had a ratio of 1:3.8 (Zhao *et al*., 2017) (Table 1). The suggestion was that *P. somniferum* does not follow a biased fractionation pattern.

**Table 1.**
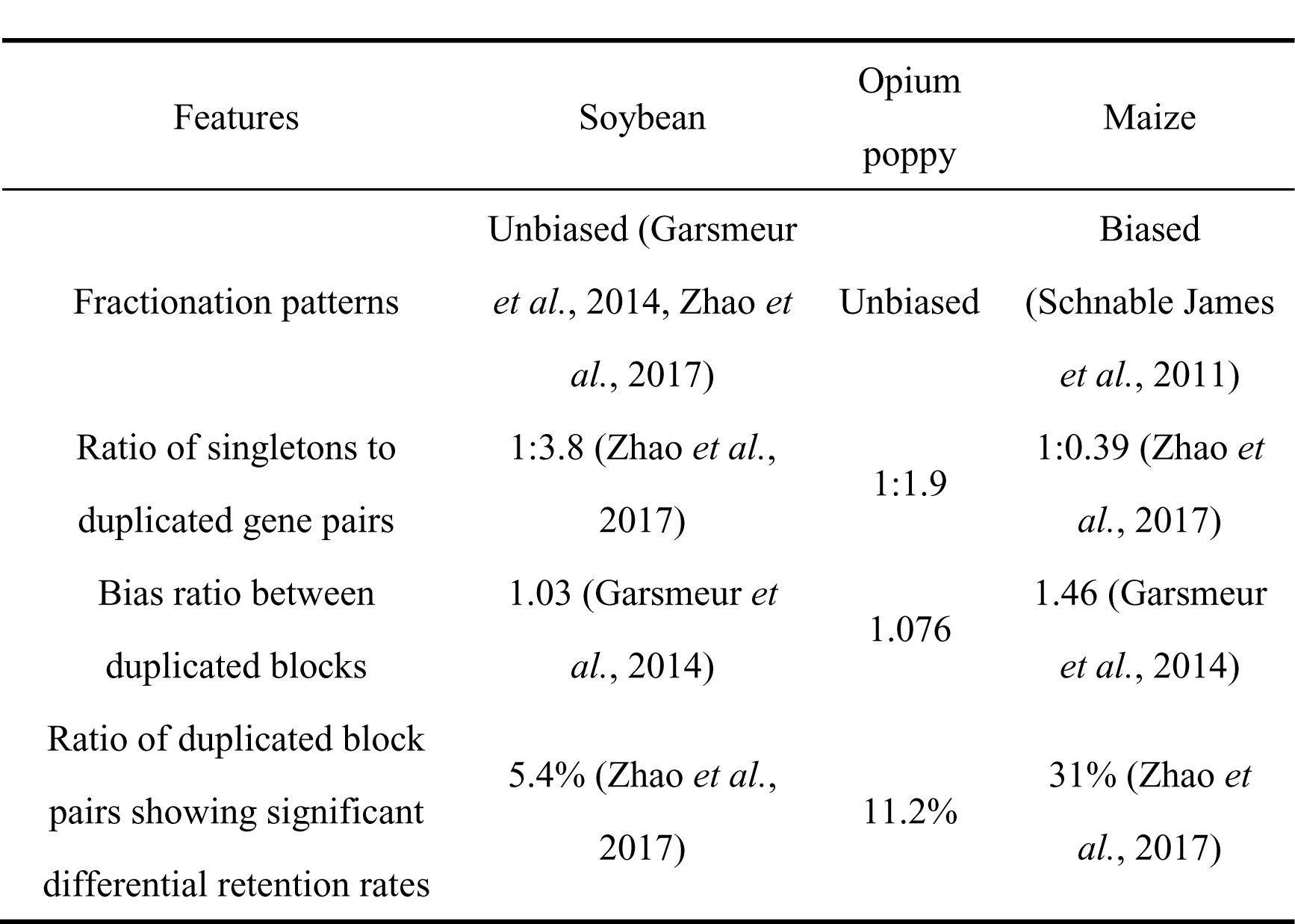
Comparisons of different subgenome features among opium poppy, soybean, and maize.

Three methods were employed to evaluate the evolutionary fractionation pattern and determine whether gene loss occurred uniformly across duplicated genomic regions. The first approach involved measuring the angle of syntenic diagonals and taking the median ratio of (longer-of-length-or-width)/(smaller-of-length-or-width), as described by (Garsmeur *et al*., 2014). A total of 170 duplicated blocks generated during the recent WGD event in *P. somniferum* were identified and then filtered to exclude blocks containing fewer genes, leaving 89 larger blocks. These blocks were used as input to estimate the fractionation pattern. The median ratio was found to be 1.08 in *P. somniferum*, between 1.46 in maize and 1.03 in soybean, which was also lower than the classical biased threshold ratio (1.10) (Garsmeur *et al*., 2014) (Table 1). In the second method, statistical tests were conducted on each pair of duplicated blocks identified in the first method to determine the fractionation pattern (Zhao *et al*., 2017). Out of 89 duplicated block pairs, only 10 (11.2%) exhibited significantly different rates of gene retention (p-value < 0.05, Fisher’s exact test), a significantly lower percentage than observed in maize (31%) (p-value < 0.05, Fisher’s exact test) (Zhao *et al*., 2017) (Table 1). The third method aimed to compare the gene retention rates of two subgenomes of *P. somniferum* by reconstructing them and anchoring them along the seven chromosomes of *P. rhoeas*. Two different methods were used to reconstruct the subgenomes to obtain accurate results. The first method was based on k-mer differences using subgenome-specific k-mers, which is a commonly used method to phase allopolyploid as two diploid progenitors that underwent independent evolutionary pathways and accumulated specific genomic characteristics before hybridization (Jia *et al*., 2022). The second method relied on counting differential genes retained in each block (Zhao *et al*., 2017). In more detail, subphaser was used to search for subgenome-specific k-mers (**Figure S1, Table S1**), which were then combined with identified syntenic blocks to separate duplicated blocks into different subgenome sets. If we treat the k-mer differences method as the gold standard, we found that simply separating the duplicated block pairs into group1 and group2 blocks based on the differences in retention rates could also have nearly 80% identity at gene level (Figure 1a, **Table S2 and Table S3**), resulting in 2,399 genes being inconsistently assigned to subgenome1 and 2,330 genes being inconsistently assigned to subgenome2 (**Table S4**). In conclusion, regardless of the method used, the analysis revealed that the seven reconstructed subgenomes of the *P. somniferum* chromosomes did not exhibit significantly biased fractionation pattern (Figure 1b, **Figure S2 and Figure S3**). However, it should be noted that the gene retention rates for chromosomes 1, 3, and 4 reconstructed using the second method exhibited significant bias (p-value < 0.05, Fisher’s exact test), while those reconstructed by the first method did not (**Table S3**). This discrepancy may be attributed to the fact that these three chromosomes had a higher number of duplicated blocks (about 51.8%) which would allow more syntenic genomic fragments to accumulate differences between the two subgenomes in accordance with the principle of the retained rate differences method. Taken together, these results suggest that the fractionation of the two *P. somniferum* subgenomes is unbiased.

**Figure 1.**
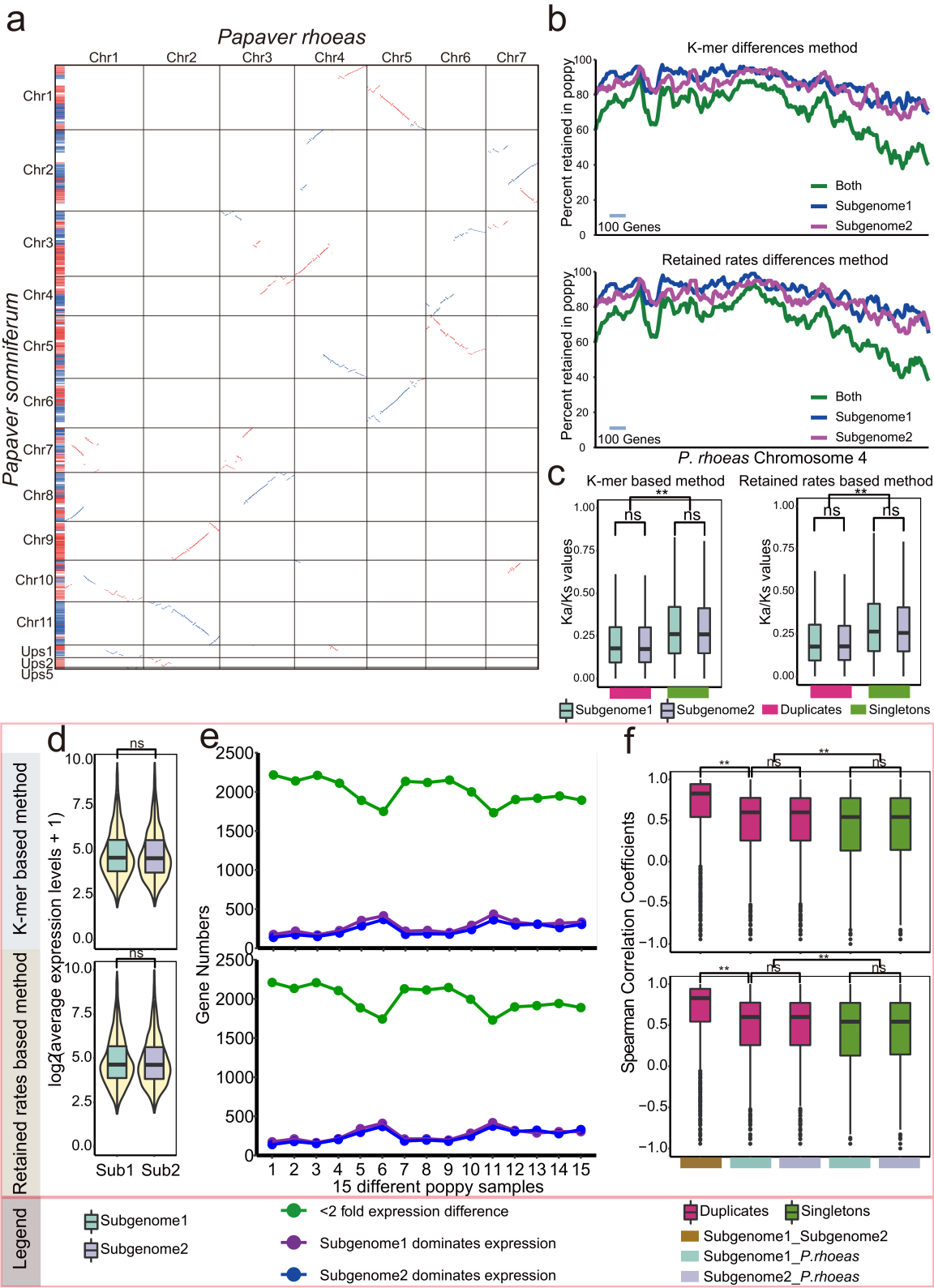
Evolutionary patterns and features between the two subgenomes of *P. somniferum* with gene-level analysis. (**a**) Dot plot analysis of interspecies syntenic blocks between *P. rhoeas* and *P. somniferum*. Red and blue dots represent genes belonging to *P. somniferum* subgenome1 and subgenome2, respectively, under the k-mer differences method. The different colors of the Y-axis indicate different subgenomes classified under the retained rates differences method. Red and blue lines indicate genes belonging to *P. somniferum* subgenome1 and subgenome2, respectively. (**b**) Genes retained percentage along *P. rhoeas* chromosome 4. The percentage (Y-axis) indicates the number of genes retained for subgenome1 (blue), subgenome2 (red) and both subgenomes (green, genes were retained both in subgenome1 and subgenome2) based on 100 genes sliding windows along *P. rhoeas* chromosome 4 (X-axis). (**c**) Comparison of the selective strength (Ka/Ks) of two types of genes (duplicates and singletons) between subgenome1 and subgenome2. (**d**) Average expression values of duplicates between subgenome1 and subgenome2 using expressed genes (TPM >= 1). (**e**) Comparison of gene expression patterns of duplicates between subgenome1 and subgenome2 using expressed genes. Domination indicates a larger than 2-fold expression difference. (**f**) Comparison of the Spearman correlation coefficients of two types of genes (duplicates and singletons) between subgenome1 and subgenome2 across six different tissues. The outlier dots in c and d were masked by ggplot2. The selective strength and Spearman correlation coefficient were calculated via pairwise comparison of *P. somniferum* genes, and their orthologs in *P. rhoeas*. The statistical analysis was conducted via the Wilcoxon test. “ns” indicates not significant (p-value > 0.05). Asterisk ** indicates adjusted p-value < 0.05 and * indicates p-value < 0.05. P-value < 0.05 was considered as significant, while adjusted p-value < 0.05 was considered as significant under the Benjamini & Hochberg correction.

Subgenome dominance refers to a bias toward higher expression from one subgenome compared to the other subgenome. Numerous studies have demonstrated a correlation between the fractionation model and subgenome dominance (Garsmeur *et al*., 2014, Zhao *et al*., 2017, Cheng *et al*., 2018, Alger and Edger, 2020). Further analysis was conducted to determine whether the association between fractionation and subgenome dominance applies to *P. somniferum*. Briefly, high-confidence genes were used as defined before to compare sequence substitution rates of duplicates and singletons relative to their orthologous genes in *P. rhoeas* (Figure 1c), gene expression patterns of duplicates among 15 different samples (Figure 1d **and** Figure 1e) and Spearman correlation coefficients between genes in each *P. somniferum* subgenome and their orthologous genes in *P. rhoeas* (Figure 1f). None of these properties showed significant divergence between the two *P. somniferum* subgenomes (adjusted p-value > 0.05, Wilcoxon test), indicating that there was no subgenome dominance observed after the recent WGD event.

However, singletons exhibited more relaxed purifying selection and higher sequence substitution rates compared to duplicates within each subgenome in the global unbiased background (Figure 1c). Similarity, gene expression correlation coefficients indicated that singletons had significantly lower expression similarity than duplicates (Figure 1f), suggesting a biased evolutionary pattern between the two groups within each subgenome (adjusted p-value < 0.05, Wilcoxon test). Moreover, this biased model was associated with specific functional features, such as lower overall expression levels and expression breadth for singletons compared to duplicates (adjusted p-value < 0.05, Wilcoxon test) (**Figure S4**). These findings suggest a connection among coding regions variations, gene expression patterns, and the genome fractionation patterns (biased or unbiased gene retentions) in *P. somniferum*.

The gene-level analysis demonstrated an unbiased fractionation model for subgenome1 and subgenome2, as well as a biased evolutionary pattern for duplicates and singletons within each subgenome, as revealed by the reconstructed subgenomes. In addition, since the two subgenomes displayed a similar skeleton and properties, only the results of the k-mer-based reconstructed subgenomes were presented in the subsequent sections, unless explicitly mentioned otherwise.

### Retention length of CNEs between two subgenomes is correlated with genome fractionation pattern

The coding and non-coding regions undergo simultaneous duplication after the WGD event (Liang and Schnable, 2018). Herein, we investigated how fractionation patterns at the gene level affect the lengths of conserved non-coding elements (CNEs), which are presumed regulatory elements. To achieve this, we utilized genes as anchors and subgenomes as scaffolds to systematically identify CNEs through pairwise comparison of non-coding sequences around *P. somniferum* genes and their orthologs in *P. rhoeas* (Turco et al., 2013).

Using the CNS Discovery Pipeline v.3.0, we identified a total of 328,129 CNEs (**Table S5**) with a length range of 15 bp - 2,631 bp, resulting in a CNEs space of 41.69 Mbp (Turco *et al*., 2013). Among these, 90% had a length ranging from 15 bp to 287 bp (Figure 2e, **Table S6**). Furthermore, to verify the consistency of identified CNEs, we used another independent CNEs discovery pipeline, the dCNS software, which generated 226.84 Mbp of CNEs space (**Table S7**) (Song *et al*., 2021). This value was larger than the previously generated CNEs space (Turco *et al*., 2013), potentially due to dCNS software used all homologous genes as anchors, while the CNS Discovery Pipeline only used high-confidence genes as input. Also, dCNS software adopted a 100 kb range to find CNEs, while CNS Discovery Pipeline adopted a 10 kb range. Although dCNS software and CNS Discovery Pipeline had differences in their CNEs discovery methods, they showed a high level of consistency in identifying CNEs, with about 90% of the CNEs identified by CNS Discovery Pipeline being present in the CNEs space generated by dCNS software. Therefore, the CNEs space generated by CNS Discovery Pipeline was used for subsequent analysis.

**Figure 2.**
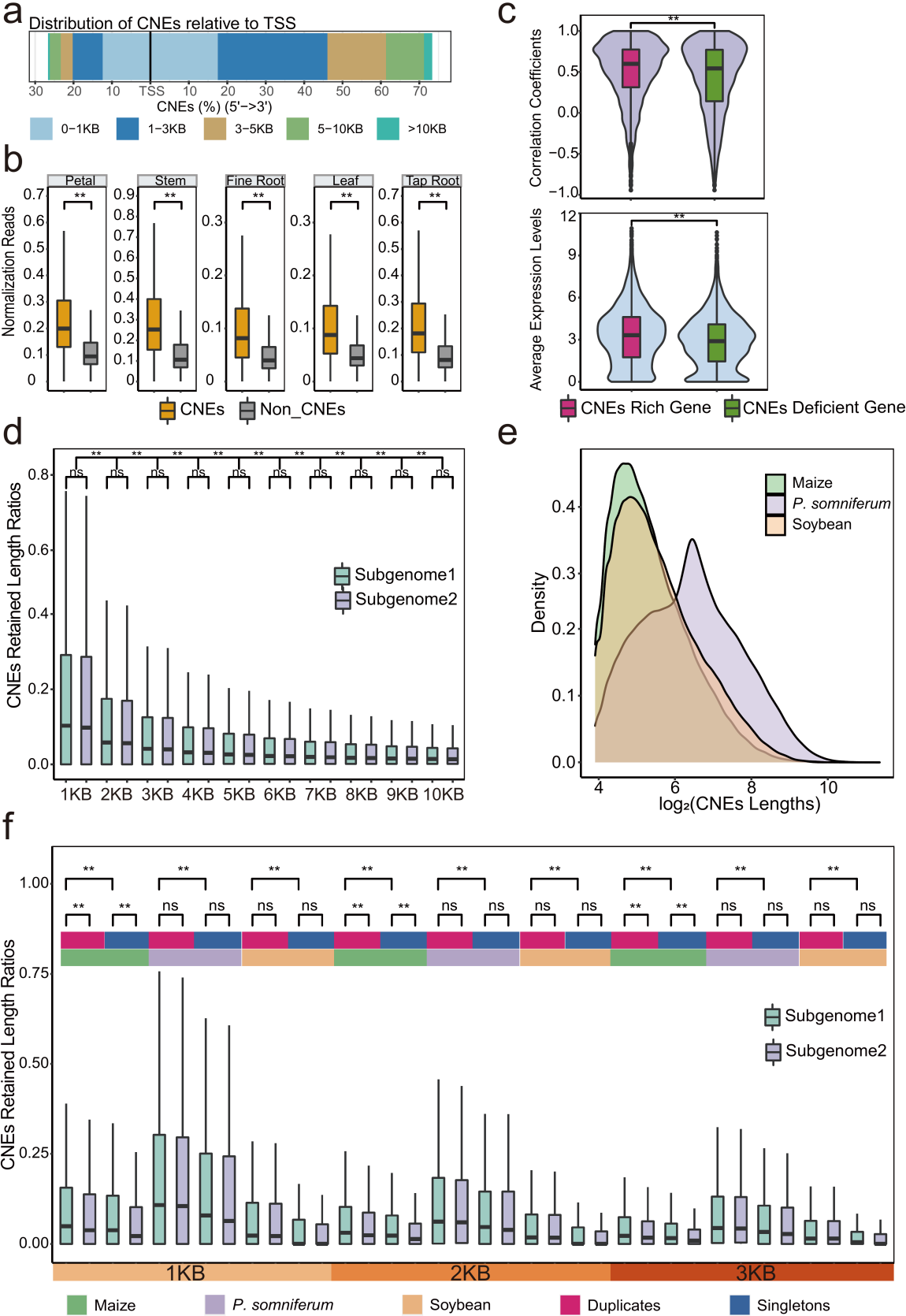
Properties and evolutionary trajectory of CNEs. (**a**) The distribution of *P. somniferum* CNEs relative to the transcription start site (the distance from the CNEs to the TSS of the nearest gene). (**b**) Comparison of normalised overlapped ATAC-seq reads between CNEs and Non_CNEs within 1 kb upstream of TSS. Sequence longer than 30 bp were used for analysis. Normalised overlapped ATAC-seq reads were calculated by the number of overlapped ATAC-seq reads divided by the length of each CNEs or Non_CNEs. (**c**) Comparison of the Spearman correlation coefficients and average expression levels between CNEs_rich genes and CNEs_deficient genes. Spearman correlation coefficients were calculated via pairwise comparison of *P. somniferum* genes and they were orthologous in *P. rhoeas*. Average expression values were calculated based on the mean of RNA-seq data across six different tissues, then transformed to log2 expression values. (**d**) Comparison of CNEs retention ratios between subgenome1 and subgenome2 for 10 different regions. For example, 2 kb represents CNEs retained ratios within 2 kb upstream regions of their related gene. (**e**) The length distribution of CNEs for three different species. (**f**) Comparison of two types of genes (duplicates and singletons) related to CNEs retained length ratios between subgenome1 and subgenom2 for three different species within 3 kb upstream regions. The outlier dots in b, d, and f were masked by ggplot2. Note: The specific comparison line directly located at the top of a specific barplot indicates that this barplot was used for statistical comparisons, like the bottom comparison lines in d. The specific comparison line located at the top of the medium of two barplots indicates that the two barplots were first merged, then used for statistical comparison, like the top comparison lines in d. The statistical analysis was conducted via the Wilcoxon test. “ns” indicates not significant (p-value > 0.05). Asterisk ** indicates adjusted p-value < 0.05 and * indicates p-value < 0.05. P-value < 0.05 was considered as significant, while adjusted p-value < 0.05 was considered as significant under the Benjamini & Hochberg correction.

The analysis of the location of CNEs relative to genes showed that 29.99% of CNEs (number) were located within 1 kb regions of the transcription start site (TSS) (Figure 2a). Moreover, these CNEs contained more normalised ATAC-seq reads within 1 kb regions upstream of TSS (Figure 2b) and intronic regions (**Figure S5**) compared to non-conserved non-coding elements (Non_CNEs) (adjusted p-value < 0.05, Wilcoxon test). In addition, within the 1 kb regions upstream of the TSS, CNEs significantly overlapped (p-value < 0.05, Fisher’s exact test) with putative TFBSs predicted by PlantTFDB (Jin *et al*., 2017, Tian *et al*., 2020) and ACRs (**Table S8, Figure S6**). These observations suggest that CNEs have potential regulatory functions at the genetic and epigenetic levels. Furthermore, CNEs were found to have regulatory functions at the transcription level. CNEs_rich genes and CNEs_deficient genes were separated based on the first, and the fourth quantile of CNEs retained lengths, respectively. CNEs_rich genes had higher expression correlation coefficients with orthologous genes in *P. rhoeas* and had higher average expression values than CNEs_deficient genes, partly indicating the impact of CNEs variations on transcription (adjusted p-value < 0.05, Wilcoxon test) (Figure 2c). Besides, “transcription factor activity” (GO:0003700, p-value < 1e-12) was significantly enriched in CNEs_rich genes, consistent with previous studies (Schnable *et al*., 2011, Burgess and Freeling, 2014).

We also explored the evolutionary fate of CNEs following WGD. The CNEs of *P. somniferum* were assigned into two classes based on their relative genes, namely subgenome1 and subgenome2-related CNEs. The retained length ratios of subgenome1 and subgenome2-related CNEs within 10 different upstream regions showed no significant difference, suggesting that an unbiased fractionation model also worked in the non-coding regions (p-value > 0.05, Wilcoxon test) (Figure 2d). This observation was further supported by comparing the retained length ratios for each chromosome (p-value > 0.05, Wilcoxon test) (**Figure S7**). Also, the relative retained numbers of CNEs between subgenome1 and subgenome2 were compared, and no significant difference was detected, supporting the above results (p-value > 0.05, Wilcoxon test) (**Figure S8**). Moreover, a comparison of different regions for CNEs retained length proportions showed that the proportion of CNEs decreased with the distance of CNEs to their related genes, with a higher proportion of sequences retained as CNEs in regions closer to the genes (adjusted p-value < 0.05, Wilcoxon test). This observation may be due to the stronger effect of purifying selection in these regions closer to the genes (Figure 2d). Additionally, considering the intrinsic properties of different species, the timing of WGD events, and distinctive genomic fractionation patterns may play the key role between two subgenomes, we included maize and soybean in our analysis. Although WGD events in maize and soybean occurred at approximately the same time (∼13 Mya, prior to the most recent WGD time in *P. somniferum*), the evolutionary fates are distinct, with maize exhibiting biased fractionation and soybean exhibiting unbiased fractionation (Zhao *et al*., 2017). CNEs in maize and soybean were identified using the same pipeline, based on their subgenomes constructed in previous studies (Zhao *et al*., 2017, Brohammer *et al*., 2018). The median lengths of CNEs in maize and soybean were significantly lower (36 and 41, respectively) than in *P. somniferum* (84), possibly due to differences in the timing of WGD (Figure 2e, **Table S6**). In addition, CNEs retained length ratios showed no significant divergence between the two subgenomes in soybean (adjusted p-value > 0.05, Wilcoxon test) but were divergent in maize subgenomes (adjusted p-value < 0.05, Wilcoxon test) (Figure 2f), consistent with previous findings (Figure 2d). Furthermore, evolutionary patterns were divergent between duplicates and singletons-related CNEs in each species (adjusted p-value < 0.05, Wilcoxon test) (Figure 2f), suggesting that intrinsic properties differ between duplicates and singletons at both gene and non-coding levels. Overall, these results suggest that the evolutionary trajectory of CNEs (presumed regulatory sequences) retained ratios between subgenomes is associated with the fractionation pattern of the genome at the gene-level.

### Complex associations among coding regions variations, CNEs retained lengths, gene expression profiles and the status of duplicates or singletons

Previous studies have highlighted the significance of divergence in gene expression patterns and regulatory systems in shaping phenotypic changes among closely related species, compared to variation in coding DNA sequences (King and Wilson, 1975, Romero *et al*., 2012, Yang and Wang, 2013, Li *et al*., 2020a). Recent research using deep neural network models and high-throughput experiments has demonstrated the high evolvability of regulatory elements and their impact on gene expression (Vaishnav *et al*., 2022). However, divergence in gene expression patterns could also be induced by, or correlated with, variation in gene coding regions, as shown in both natural and experimental populations (Li *et al*., 2005, Shen *et al*., 2022). Herein, results suggest the interplay among coding regions variations, CNEs retained lengths, gene expression profiles, and the status of duplicates or singletons is complex (Figure 1c, Figure 1f **and** Figure 2c, **Figure S4**). Therefore, this study aimed to address the following questions: (1) which evolves faster following WGD, gene coding sequence or gene expression patterns (using the relative number of divergent genes as the indicator)? Evolving faster indicates that more genes are distinctive to their ancestry copy. (2) What are the properties of transcriptomic evolutionary processes? (3) What is the impact of CNEs presence-absence variations on transcriptomic evolution? By addressing these questions, this study aimed to gain a better understanding of the mechanisms underlying transcriptomic evolution in response to WGD events.

Conservation and specialization are two important types of gene expression patterns or gene function evolution. Conservation means a gene maintains a similar expression pattern or function to that of the ancestral gene, while specialization indicates that a gene has a distinctive pattern (Assis and Bachtrog, 2013, Wang *et al*., 2016). Our initial focus was on assessing the rates of divergence in gene expression patterns. Pearson correlation coefficients (PCCs) were calculated via pairwise comparison of gene expression profiles of *P. somniferum* genes and their orthologs in *P. rhoeas.* Evolutionary trajectories for duplicates and singletons were then defined to quantitatively evaluate the transcriptomic evolution following WGD. For duplicates, two genes were retained following WGD, and we calculated PCCs between the expression profiles of subgenome1 duplicate copy (D1) and the *P. rhoeas* ancestral copy (A) (R_D1,A_) and between the expression profiles of subgenome2 duplicate copy (D2) and the *P. rhoeas* ancestral copy (A) (R_D2,A_). The same method was used for singletons. The correlations for subgenome1 singletons (S1) with *P. rhoeas* ancestral copy (A) (R_S1,A_) and the correlation for subgenome2 singletons (S2) with *P. rhoeas* ancestral copy (A) (R_S2,A_) were also calculated. We defined the baseline for expression divergence (R_O1,O2_) as the mean of PCCs for single-copy orthologous genes between *P. somniferum* single-copy genes and their single-orthologs in *P. rhoeas*, which we identified using three *Papaver* species (*P. rhoeas*, *P. somniferum* and *Papaver setigerum*).

After removing low expression genes (normalised expression levels < 1), 6,944 pairs of duplicates were retained (86.1%). Presumably, R_O1,O2_ represents the expected PCCs. R_D1,A_ and R_D2,A_ were compared with R_O1,O2_ for each paralog pair (duplicates). Four types of evolutionary trajectories were defined, including both_conservation, specialization_1, specialization_2, and both_specialization (Table 2). For example, for specialization_1, the subgenome1 duplicate copy expression profile should be different from the ancestral copy, while the subgenome2 copy should be similar to the ancestral copy, resulting in R_D1,A_ < R_O1,O2_ and R_D2,A_ >= R_O1,O2_. In total, 3,837 both_conservation, 675 specialization_1, 723 specialization_2 and 1,709 both_specialization cases were identified (Table 2, **Table S9**). Approximately 55.26% of the gene pairs were retained through total transcriptional profiles conservation with the ancestral genes. For singletons, after removing low expression genes (normalised expression levels < 1), 1,599 (72.5%) and 1,464 (72.1%) of genes were retained for subgenome1 and subgenome2, respectively, indicating a higher percentage of low-expression genes compared to duplicates. Next, using the same methods as with duplicated, four evolutionary trajectory types (sub1_conservation, sub1_specialization, sub2_conservation, and sub2_specialization) were defined (Table 3). For example, for sub1_conservation indicates that only one copy with a syntenic relationship should be retained in subgenome1 after WGD, and its expression profile should be similar to that of the ancestral copy, thus showing that R_s1,A_ >= R_O1,O2_. Overall, 955 sub1_conservation, 644 sub1_specialization, 880 sub2_conservation, and 584 sub2_specialization cases were identified (Table 3, **Table S10**). About 59.72% - 60.11% of singletons had conserved expression profiles with the ancestral genes. Consistent with duplicates, most genes were retained even for singletons through transcriptional profiles conservation with the ancestral genes, (Table 2 **and** Table 3). To compare the divergence rates between gene coding sequences and gene expression patterns, we defined the evolutionary trajectories for coding sequences via pairwise comparison of selection pressures (Ka/Ks values) of *P. somniferum* genes and their orthologs in *P. rhoeas*. For duplicates, 5,726 both_conservation, 614 specialization_1, 541 specialization_2 and 1,177 both_specialization cases were identified (**Table S11**). For singletons, 1,343 sub1_conservation, 861 sub1_specialization, 1,227 sub2_conservation and 800 sub2_specialization cases were identified (**Table S11**). Overall, 64.3% of genes (duplicates and singletons) had conserved expression profiles with the ancestral genes compared with 74.6% of the genes that underwent higher levels of selection pressure for coding sequences. Additionally, we also defined the evolutionary trajectories for coding sequences via substitution numbers (Ka + Ks values) and then identified that 86.7% of the genes had lower coding substitutions. These differences suggest that divergence in gene expression patterns is faster than divergence in gene coding sequences, especially for duplicates (p-value < 0.05, Fisher’s exact test).

**Table 2.** Classification of evolutionary trajectories of gene expression between *P. somniferum* duplicates and *P. rhoeas* ancestral copy by measuring the Pearson correlation coefficients.

| Classification | $R_{D1,A}$ | $R_{D2,A}$ | Pairs (Gene numbers) |
| --- | --- | --- | --- |
| Both_Conservation | $\geq R_{O1,O2}$ | $\geq R_{O1,O2}$ | 3,837 (7,674) |
| Specialization_1 | $< R_{O1,O2}$ | $\geq R_{O1,O2}$ | 675 (1,350) |
| Specialization_2 | $\geq R_{O1,O2}$ | $< R_{O1,O2}$ | 723 (1,446) |
| Both_Specialization | $< R_{O1,O2}$ | $< R_{O1,O2}$ | 1,709 (3,418) |
| Total | - | - | 6,944 (13,888) |

Note: $R_{O1,O2}$ is the mean Pearson correlation coefficients (PCCs) for single-copy orthologous genes identified across three *Papaver* species. $R_{D1,A}$ represents PCC between subgenome1 duplicate copy (D1) in *P. somniferum* and the *P. rhoeas* ancestral copy (A). $R_{D2,A}$ represents PCC between subgenome2 duplicate copy (D2) in *P. somniferum* and the *P. rhoeas* ancestral copy (A).

**Table 3.**
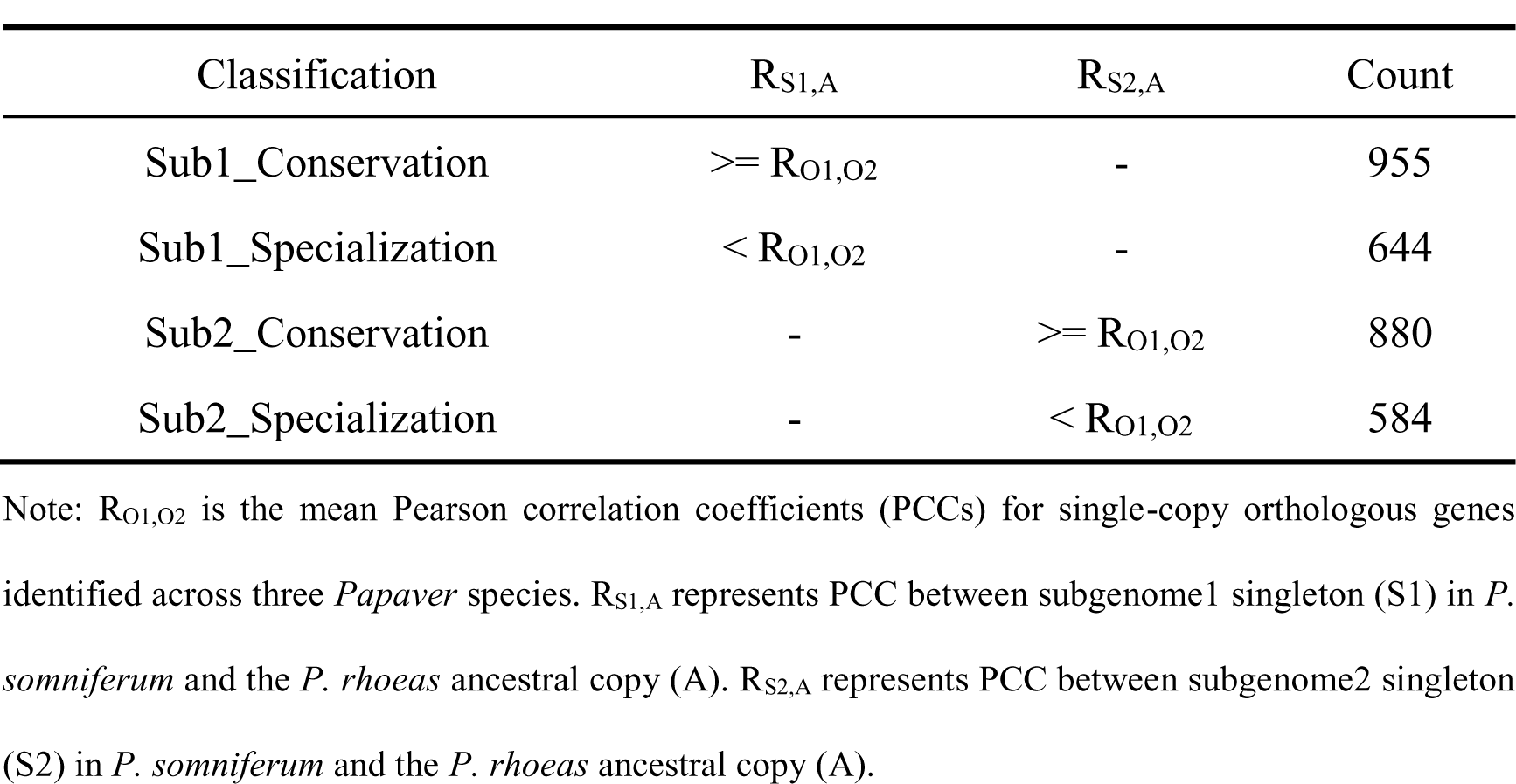
Classification of evolutionary trajectories of gene expression between *P. somniferum* singletons and *P. rhoeas* ancestral copy by measuring the Pearson correlation coefficients.

| Classification | $R_{S1,A}$ | $R_{S2,A}$ | Count |
| --- | --- | --- | --- |
| Sub1_Conservation | $\geq R_{O1,O2}$ | - | 955 |
| Sub1_Specialization | $< R_{O1,O2}$ | - | 644 |
| Sub2_Conservation | - | $\geq R_{O1,O2}$ | 880 |
| Sub2_Specialization | - | $< R_{O1,O2}$ | 584 |

Note: $R_{O1,O2}$ is the mean Pearson correlation coefficients (PCCs) for single-copy orthologous genes identified across three *Papaver* species. $R_{S1,A}$ represents PCC between subgenome1 singleton (S1) in *P. somniferum* and the *P. rhoeas* ancestral copy (A). $R_{S2,A}$ represents PCC between subgenome2 singleton (S2) in *P. somniferum* and the *P. rhoeas* ancestral copy (A).

The properties of different evolutionary process-related genes, as defined by their similarity in transcriptional profiles, were also explored. Initially, the focus was directed towards duplicates, revealing that such genes exhibited distinct characteristics. Functionally, metabolism and transport-related genes were more likely to undergo divergence in transcriptional profiles with ancestral genes. Conversely, genes associated with fundamental biological processes, such as ribosome and messenger RNA biogenesis, were conserved (Figure 3a, **Table S12**). With regards to selection pressures, coding regions for both_conservation-related genes were under greater selection pressures than both_specialization-related genes (adjusted p-value < 0.05, Wilcoxon test). However, there were no significant differences between subgenome1 and subgenome2 for specialization_1 and specialization_2-related genes (adjusted p-value > 0.05, Wilcoxon paired test) (Figure 3b, **Figure S9**). In other words, specialization_1 and specialization_2-related genes experienced transcriptional divergence between the two subgenomes, while coding regions were similar. These findings suggest that variations in coding sequences may not be the main factor contributing to the divergence in expression patterns of these genes. However, CNEs retained length ratios were significantly correlated with transcriptional profiles divergence, especially CNEs retained ratios within 1 kb regions (adjusted p-values < 0.05, Wilcoxon paired test) (Figure 3c **and Figure S9**), suggesting that CNEs variations may have a greater influence on gene expression variations. Furthermore, for singletons, conservation-related genes were under higher selection pressures in both coding and non-coding regions than specialization-related genes (adjusted p-value < 0.05, Wilcoxon test) (**Figure S10**). The functional enriched terms for conservation-related genes and specialization-related genes were consistent between singletons and duplicates (Figure 3a, **Table S13**).

**Figure 3.**
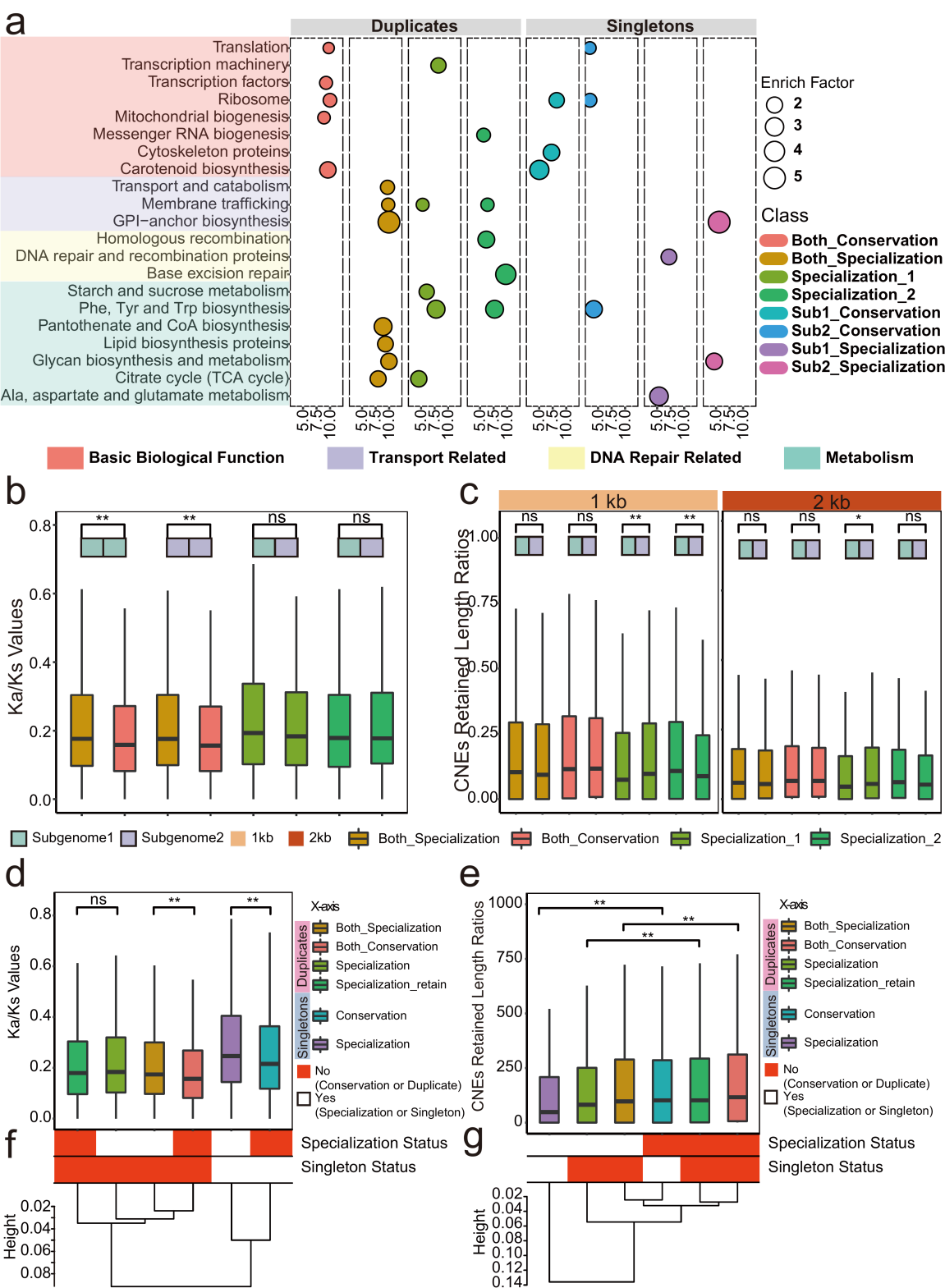
Comparison of selection pressures on coding regions and CNEs for two types of genes (duplicates and singletons) classified in different transcriptome evolutionary processes. (**a**) Enrichment analysis (KEGG) of each evolutionary process-related genes. (**b**) Comparison of the Ka/Ks values for different evolutionary processes of duplicates. (**c**) Comparison of the CNEs retained ratios for different evolutionary processes of duplicates within 2 kb regions. Comparison of the (**d**) Ka/Ks values and (**e**) CNEs retained lengths (1 kb) for different evolutionary processes between duplicates and singletons. For duplicates, genes of Specialization_1 process from subgenome1 and genes of Specialization_2 process from subgenome2 were merged into Specialization, whereas genes of Specialization_1 process from subgenome2 and genes of Specialization_2 process from subgenome1 were merged into Specialization_retain. For singletons, sub1_conservation and sub2_conservation were merged into Conservation. Sub1_specialization and sub2_specialization were merged into Specialization. The topology of cluster tree of the (**f**) Ka/Ks values and (**g**) CNEs retained lengths (1 kb) for different evolutionary processes between duplicates and singletons. The outlier dots in b, c, d, and e were masked by ggplot2. B and c share a common set of labels. The statistical analysis was conducted via the Wilcoxon test or Wilcoxon paired test. “ns” indicates not significant (p-value > 0.05). Asterisk ** indicates adjusted p-value < 0.05 and * indicates p-value < 0.05. P-value < 0.05 was considered as significant, while adjusted p-value < 0.05 was considered as significant under the Benjamini & Hochberg correction.

To systematically explore the underlying complex relationships, the results of duplicates and singletons were combined. Surprisingly, it was found that all evolutionary processes for singletons were under more relaxed selection pressures on coding regions than duplicates (Figure 3d), while CNEs variations were not significant (Figure 3e). To gain more quantitative results, we cut the distributions of Ka/Ks values and CNEs retained lengths of genes involved in different evolutionary processes (Figure 3d **and** Figure 3e) into hundreds of bins, and then counted the number of points in each bin and normalised them to the same range (ranging from 0 to 1, with larger values in a bin meaning more points located in that bin). The distance between each distribution was calculated using the bin-related profiles, then clustered. The topology of the cluster tree for selection pressure was correlated with singleton status, which was defined as duplicated genes originating from a WGD event, retained as a duplicate or singleton (Figure 3f). In addition, cluster tree topology for CNEs retained length was more closely associated with transcriptome evolutionary processes (conserved or specialized expression profiles) than with singleton status (Figure 3g). To validate the robustness of our analysis, we applied different thresholds and methods. The median of PCCs for single-copy orthologous genes across three *Papaver* species was selected as the baseline for expression divergence (stricter threshold, R_O1,O2_ = 0.647 adopted by median > 0.514 adopted by mean), and similar findings were found (**Figure S11**). Although the Spearman correlation is less powerful than the Pearson correlation, it can also be accepted and used to measure expression divergence (Casneuf *et al*., 2006, Metzger *et al*., 2017, El Taher *et al*., 2021). Therefore, a similar analysis was conducted using Spearman correlation coefficients (SCCs), yielding similar results (**Figure S12**). Our results systematically and deeply compared multi-features of the genomic and transcriptomic variations and revealed complex interplays among coding sequence variations, CNEs retained ratios and transcriptomic divergence following WGD for duplicates and singletons.

Previous findings showed that variable CNEs are associated with accessible chromatin regions (ACRs) (Yocca *et al*., 2021). The normalised ATAC-seq reads within 1 kb regions upstream of TSS were compared between conserved and specialized genes in the lists of specialization_1 and specialization_2 (their *P. rhoeas* ancestral copies must be tissue-specific) to comprehensively assess the above relationship and test whether part of the impact of CNEs variations on expression differences might be associated with the gain or loss of ACRs (Table 2, **Table S9**). The selective standards were applied due to these duplicates may have already undergone transcriptional divergence, and the tissue-specific ancestral copies have the simplest expression profiles, making the conserved genes have similar tissue-specificity or high expression levels in those tissues. In contrast, the specialized genes may not have such tissue specificity or high expression levels compared to their *P. rhoeas* ancestral copies. Therefore, if CNEs variations are associated with ACRs variations that influence transcriptional divergence, a significantly different number of normalised ATAC-seq reads must be contained between conserved and specialized genes for such specific tissue. For instance, if one *P. rhoeas* ancestral copy is a petal-specific gene, the conserved gene may contain more petal-related normalised ATAC-seq reads than the specialized gene.

Herein, 10,556 tissue-specific expressed genes were identified in *P. rhoeas* using Tau values (larger than 0.8) (Kryuchkova-Mostacci and Robinson-Rechavi, 2017). The previously mentioned statistical test was then performed using ATAC-seq data from four distinctive tissues of *P. somniferum*. Although only 207 tissue-specific expressed genes in *P. rhoeas* belonged to the list of specialization_1 and specialization_2, there was a significantly different number of normalised ATAC-seq reads contained between conserved genes and specialized genes (p-value = 0.0017, paired and single-sided Student’s t-test) (**Figure S13**). We demonstrated instances where CNEs variations were associated with variations in ACRs, which influenced transcriptional divergence. For example, a *P. rhoeas* homolog (*Prh07G15430.0*) of *Arabidopsis PRN2* was highly expressed in the fine root (Figure 4a) and had two orthologs in *P. somniferum* (*Pso03G09860.0* and *Pso02G15220.0*). *Pso03G09860.0* was located on subgenome1 with similar specificity to that of *Prh07G15430.0*, while *Pso02G15220.0* shifted to stem specificity (Figure 4a). However, there were no significant differences in coding regions (identity: 98.466 vs. 96.933, Ka/Ks value: 0.06 vs. 0.09). Furthermore, a core CNEs-region with a functional fine root-related ACR of nearly 300 bp was lost upstream of *Pso02G15220.0* compared with *Pso03G09860.0*, which was associated with the loss of fine root-specific ACR and the origin of a novel stem-specific ACR (Figure 4c). We hypothesized that this novel stem-specific ACR, which evolved from the lost region, might have contributed to the divergence of the *Pso02G15220.0* expression profile. To obtain more solid hypotheses and findings, we expanded our analysis to include another *Papaver* species, *Papaver setigerum* (Troy poppy). Studies have shown that *P. setigerum* underwent two rounds of WGDs following the divergence of its common ancestor from *P. rhoeas* (Yang *et al*., 2021), implying that there might be two genes in *P. somniferum* and four genes in *P. setigerum* for a single gene in *P. rhoeas*. Herein, syntenic genes between *P. setigerum* and *P. rhoeas* were identified and used as anchors to generate CNEs space. Four orthologs of *Prh07G15430.0* were retained in *P. setigerum*, of which three had similar expression profiles with *Pso03G09860.0* and *Prh07G15430.0*, and only one had similar expression profiles with *Pso02G15220.0* (Figure 4b). However, the two groups did not have significantly different coding regions (identity: 98.16, 95.918, 98.466 vs. 96.933, Ka/Ks value: 0.08, 0.20, 0.06 vs. 0.10). Additionally, the three conserved expression profiles-related genes contained the core CNEs-region, whereas the divergent gene lost this core region (Figure 4d), which is consistent with the previous hypothesis. Besides, the result showed that the tissue specificity shift for *Pso06G01300.0* might be due to upstream CNEs variations, leading to novel petal-specific ACR. This finding was also validated in the *P. setigerum* species (**Figure S14**).

**Figure 4.**
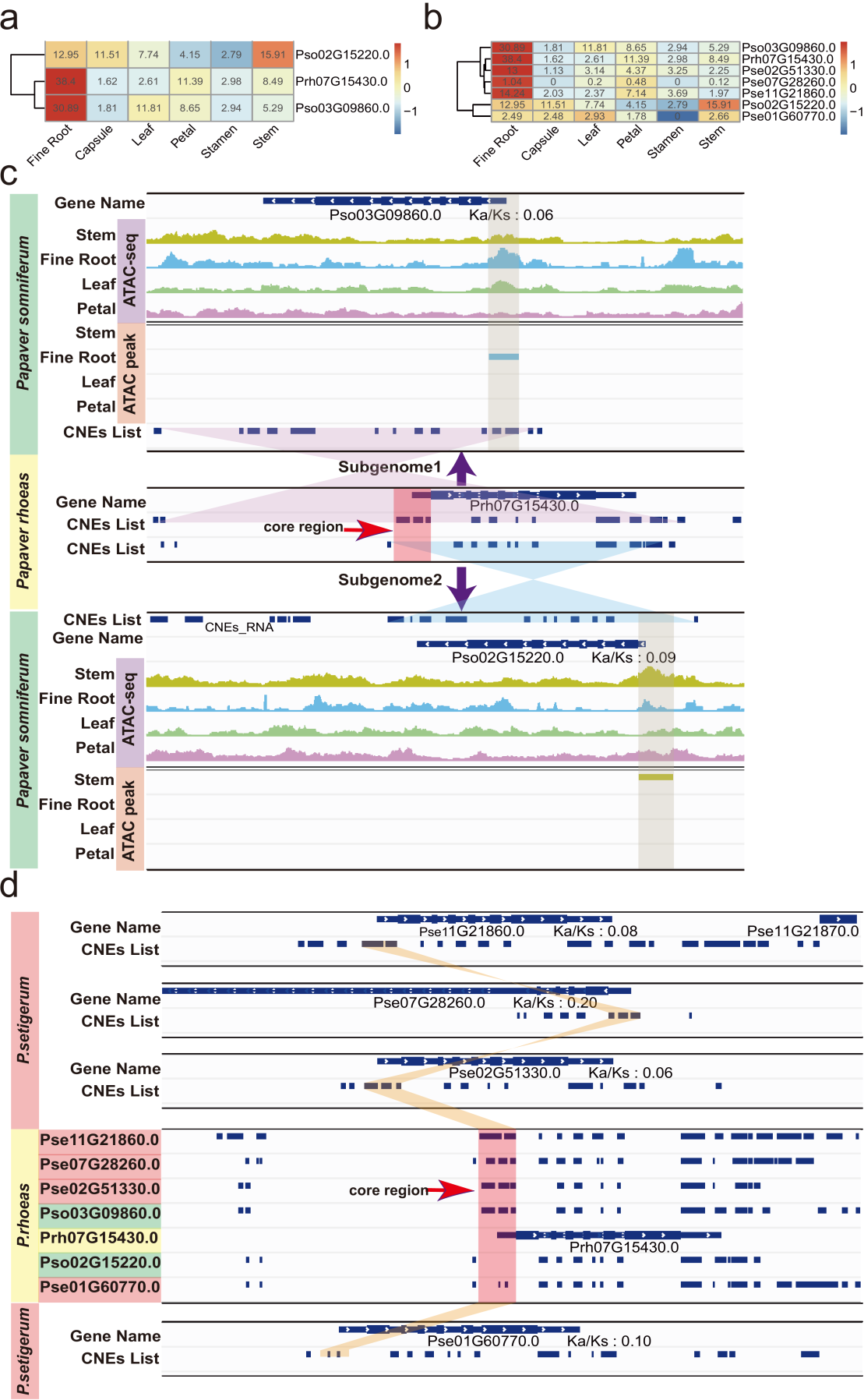
The impact of CNEs presence-absence variations on transcriptomic evolution. (**a**) Heatmap of gene expression profiles for *Prh07G15430.0* and its two orthologs in *P. somniferum* (*Pso03G09860.0* and *Pso02G15220.0*). (**b**) Heatmap of gene expression profiles for *Prh07G15430.0,* its two orthologs in *P. somniferum* and its four orthologs in *P. setigerum*. An example of CNEs variations associated with the origin of novel stem-specific ACR obtained by comparing (**c**) *P. somniferum* and *P. rhoeas*, (**d**) *P. setigerum* and *P. rhoeas*. The brown transparent boxes represent ACRs. The pink and blue transparent boxes represent CNEs regions for subgenome1 and subgenome2, respectively. The red transparent boxes represent the core CNEs-region. The orange transparent boxes represent the core CNEs-region variations across four orthologs in *P. setigerum*.

In addition, as inspired by previous research that suggests positional variations of CNEs play a role in promoting plant evolution, we identified an example of gene expression differences that may be caused by positional variations of CNEs in *P. somniferum*. *Prh05G21390.0* (**Figure S15a**), encoding a *P. rhoeas* homolog of *Arabidopsis* belonging to filament-like plant proteins, which are involved in microtubule binding and secondary cell wall deposition (Oda *et al*., 2015, Chen *et al*., 2016), had stem specificity. *Pso01G37190.0* and *Pso06G20270.0*, the two orthologs of *Prh05G21390.0,* contained a total of 4,407 bp CNEs and 1,672 bp CNEs, respectively (Ka/Ks values: 0.21 and 0.27) (**Figure S15b**). However, only *Pso06G20270.0,* with less CNEs space, maintained stem specificity. Furthermore, the CNEs of *Pso01G37190.0* contained positional variations, resulting in the lack of CNEs within the 3 kb regions upstream of *Pso01G37190.0,* which may lead to expression divergence and loss of stem specificity (**Figure S15b**). Four orthologs of *Prh05G21390.0* in *P. setigerum* were also identified, of which only one had stem specificity (**Figure S16a**). The other three orthologs lost the stem specificity and had more similar expression profiles to that of *Pso01G37190.0* (**Figure S16a**). The three orthologs also contained more CNEs in total but also experienced positional variations in the core CNEs-region, similar to *Pso01G37190.0* (**Figure S16b**). These findings demonstrate that CNEs variations can lead to faster evolution of genes even in the absence of mutations in the coding regions, providing a new insight into plant phenotypic evolution.

### Certain binding motifs are enriched in CNEs for distinctive transcriptome evolutionary processes

CNEs serve as proxies of cis-elements that transcription factors can bind to regulate gene expression. Previous results have indicated that distinctive transcriptional profiles-related genes contained different biological functions and CNEs retained ratios (Figure 3a **and** Figure 3c). Moreover, some CNEs variations are associated with transcriptional divergence (Figure 4, **Figure S14-S16**), which has piqued our interests in the putative regulatory functions of theses CNEs. Therefore, we want to investigate whether certain binding motifs are enriched in these CNEs to maintain specific biological functions through regulatory factors. To this end, we analyzed both_conservation and both_specialization gene lists because they contained a relatively large number of genes compared with other evolutionary processes (Table 2 **and** Table 3) and showed significant divergence in multifarious aspects (Figure 3a-c). In addition, we only considered CNEs located 1 kb upstream of genes.

First, the two types (both_conservation and both_specialization) of CNEs annotations were randomly shuffled 10,000 times across their respective 1 kb upstream of genes to obtain two random backgrounds. Next, 484 motifs in *Arabidopsis* were then used to scan two types of CNEs and their respective 10,000 times backgrounds (O’Malley *et al*., 2016). Finally, these motifs were classified into two classes (significant and non-significant enrichment) based on 99% percentile and 1.25-fold enrichment scores. The both_conservation and both_specialization contained 132 and 155 significantly enriched motifs, respectively. In addition, 29 and 52 motifs were only enriched in both_conservation and both_specialization, respectively (Figure 5a). Most of the “only-enriched” binding motifs displayed specific biological functions (**Table S14 and Table S15**), and their global enriched patterns (adjusted p-value < 0.01) were similar to the functional features of each evolutionary process (**Figure S17,** Figure 3a). For example, for both_conservation related CNEs analysis, the annotations of these motifs are related to the basic biological functions, such as *DEL2* was associated with cell cycle, while *bHLH80* was associated with chlorophyll and photosynthesis (Soudy *et al*., 2020) (Figure 5b **and** Figure 5c, **Table S14**). However, binding motifs enriched in the both_specialization contained many specifically enriched terms associated with multiple stress responses, development processes, and secondary metabolism compared with binding motifs enriched in both_conservation (**Figure S17**), such as *MYR2 and HHO3* (associated with starvation and nutrients) (Soudy *et al*., 2020) (Figure 5d, **Figure S18, Table S15**).

**Figure 5.**
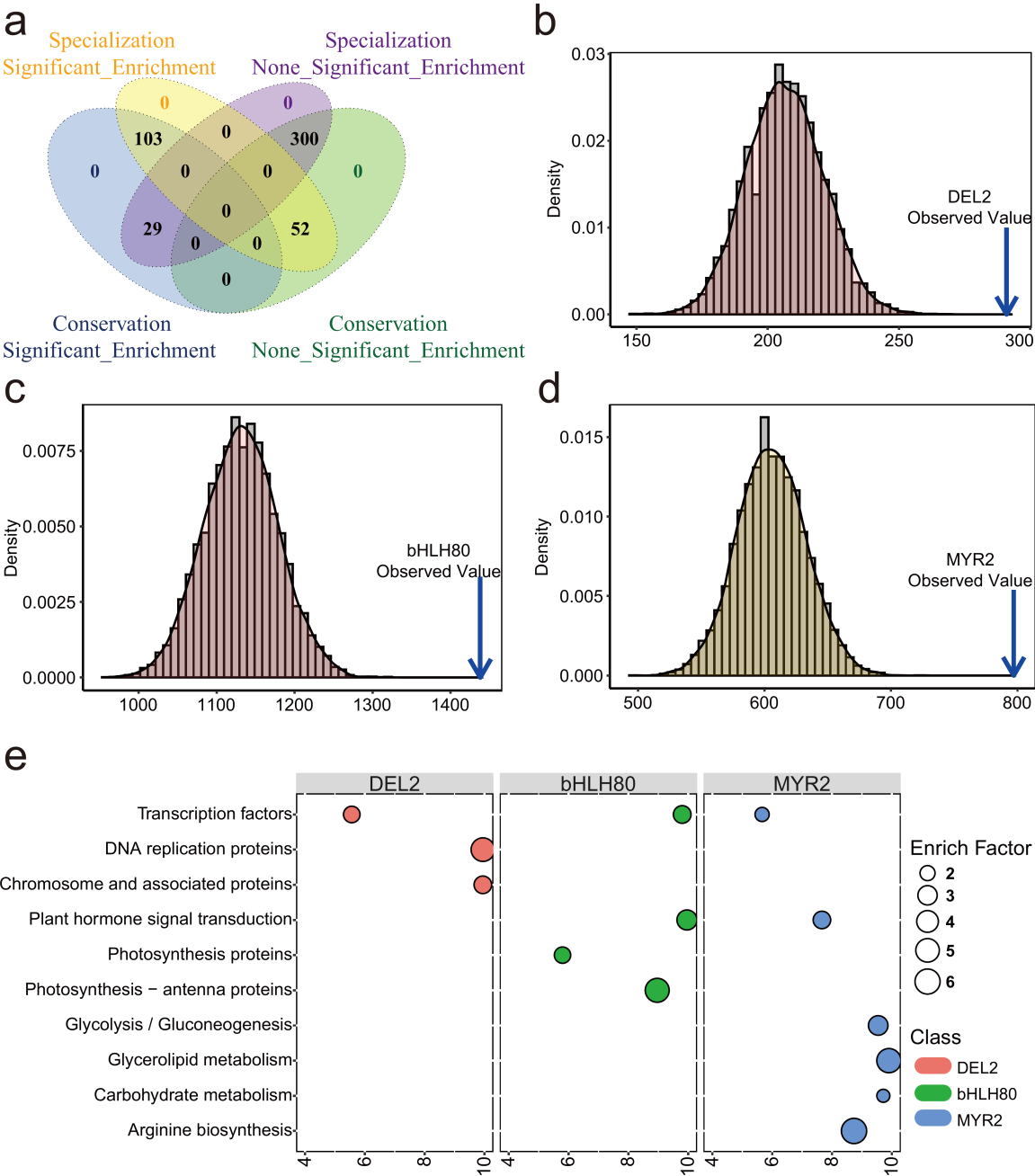
Motifs enrichment for CNEs of distinctive evolutionary processes. (**a**) Venn diagram displaying the number of different types of motifs enriched in two distinct evolutionary processes. Comparison of observed values with background distribution for (**b**) *DEL2*, (**c**) *bHLH80*, (**d**) *MYR2*. (**e**) KEGG enrichment results of *DEL2*, *bHLH80* and *MYR2* putative targeted genes.

Various functional motifs are enriched in different CNEs. Therefore, we assessed whether their putative targeted genes exhibit similar enrichment patterns. The targeted genes of *DEL2* were enriched in functional categories related to “DNA replication proteins” and “Chromosome and associated proteins”, whereas those of *bHLH80* were enriched in “Photosynthesis - antenna proteins” and “Photosynthesis proteins” (Figure 5e). In addition, for both_specialization lists, the targeted genes of *MYR2* were related to many metabolite activities and signal transduction, such as “Glycolysis / Gluconeogenesis”, “Glycerolipid metabolism” and “Arginine biosynthesis” (Figure 5e).

In brief, a collection of specific enriched binding motifs was identified. The annotations of these motifs and their associated genes showed similar functional characteristics for each evolutionary process (Figure 3a). These findings suggest that transcriptomic evolution, resulting from CNEs variations, may be associated with variations in specific regulatory elements.

### The evolutionary patterns for BIA gene cluster in non-coding regions

A trademark feature of *P. somniferum* is its ability to produce diverse range of medicinal benzylisoquinoline alkaloids (BIAs), such as morphine and noscapine. Most genes encoding the morphine and noscapine biosynthetic pathways are organised in an extreme stem specificity gene cluster (BIA gene cluster) in the *P. somniferum* genome following a recent WGD event and subsequent rearrangements (Guo *et al*., 2018). However, little attention has been paid to the regulatory mechanisms and evolutionary history underlying such tremendous innovations in expression profiles and phenotypes (Yang *et al*., 2021), which raises our interest and question how cis-elements have evolved with transcriptional evolution for the BIA gene cluster. Of the genes within the BIA gene cluster, only four genes (*PSSDR1*, *CYP719A21*, *PSMT1* and *SALR*) were located in the constructed subgenome1, and few or no CNEs were retained upstream of each gene. The upstream environment of *PSSDR1* and *SALR* were completely different from their orthologs in *P. rhoeas*, while only the *PSMT1-CYP719A21* cluster contained few CNEs, which suggested this tremendous transcriptional evolution was likely associated with rapid changes in the regulatory environment (**Table S16**).

We then evaluated why the *PSMT1-CYP719A21* cluster retained a few CNEs and what the association between these CNEs and transcriptional evolution. We identified two *PSMT1-CYP719A21* clusters in *P. rhoeas* chromosome 3 (*Prh03G43060.0-Prh03G43070.0* (cluster1) and *Prh03G45990.0-Prh03G46000.0* (cluster2)). This phenomenon may be due to the specific local duplication of the *PSMT1-CYP719A21* cluster in *P. rhoeas*. The genes homologous to the *PSMT1-CYP719A21* cluster were highly expressed in the fine roots of *P. rhoeas*, while they shifted to stem specificity in *P. somniferum* (Figure 6a). *PSMT1-CYP719A21* was arranged in a specific structure of head-to-head in opposite directions in each species, and most of the aforementioned CNEs were located in this head-to-head intergenic region (Figure 6b). Therefore, motif enrichment analysis was performed to evaluate the functions of these CNEs. The results revealed that three MYB family-related motifs, (*MYB96*, *MYB30* and *MYB15*) were enriched in the retained CNEs. Functional annotations showed that these binding motifs were associated with some root-specific functions, such as regulation of lateral root growth, defense response and response to water deprivation (Vailleau *et al*., 2002, Ding *et al*., 2009, Seo *et al*., 2009). Additionally, a stem-specific ACR was located in this specific intergenic region and overlapped with two CNEs and a root related ACR in *P. somniferum* (Figure 6b), which was also the unique stem-related ACR in the 30 kb genomic environment surrounding this gene cluster (**Figure S19**). In conclusion, we systematically assessed the regulatory regions of the *PSMT1-CYP719A21* gene cluster based on genomic, transcriptomic, and epigenetic data. This study provides evidence and speculations on the origin of the BIA gene cluster and dramatic expression shifts.

**Figure 6.**
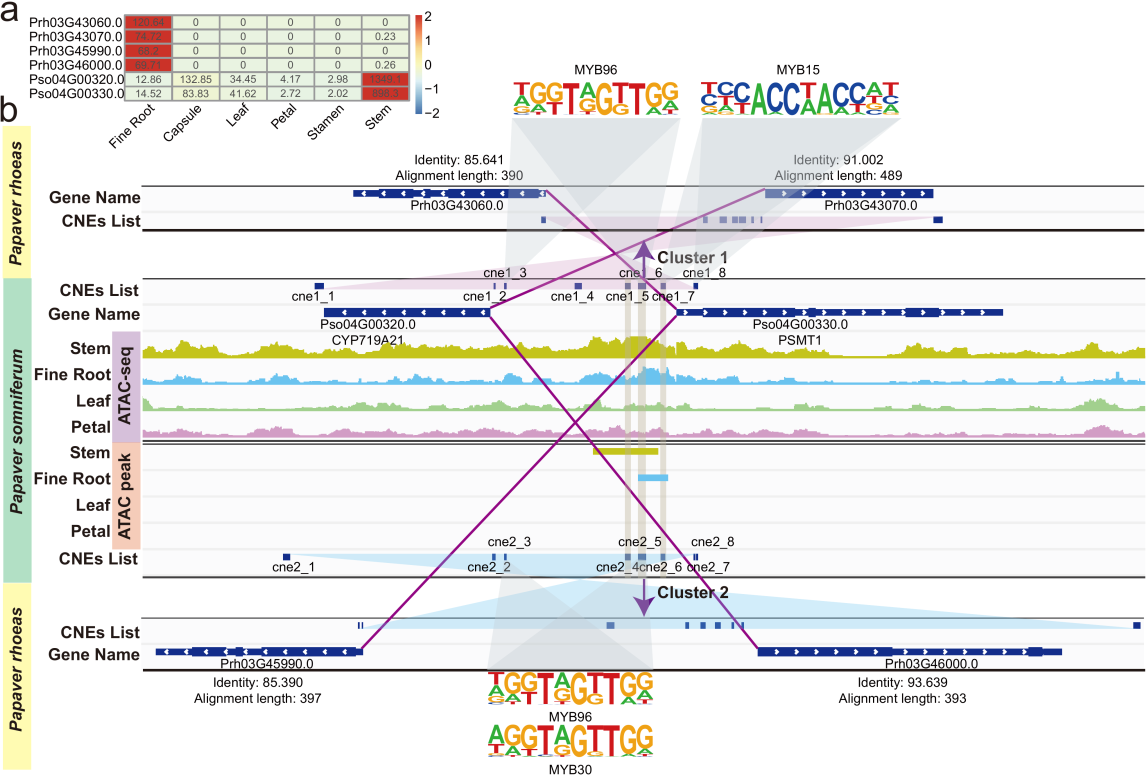
Co-regulation and co-evolution of *PSMT1-CYP719A21* cluster. (**a**) Heatmap of gene expression profiles for *PSMT1-CYP719A21* cluster genes in *P. rhoeas* and *P. somniferum*. (**b**) The origin of a novel stem-specific ACR using CNEs as scaffolds. The brown transparent boxes represent ACRs. The pink and blue transparent boxes represent CNEs regions for cluster1 and cluster2, respectively. The grey transparent boxes represent binding motifs-related CNEs. Identity and alignment length were calculated using BLAST software.

## Discussion

In this study, *P. somniferum* was used as a model system to investigate the evolutionary patterns of coding sequences, conserved non-coding elements (CNEs) and transcriptomes, respectively, and their interactions subsequent to the whole-genome duplication (WGD). Our findings provide evidence that variation in CNEs significantly affects transcriptomic evolution after WGD and may serve as an important source of evolutionary innovation in plants.

### The fractionation model plays a synergistic role in both coding and non-coding regions

Since the assembly of the first maize subgenomes in 2011 and the subsequent characterization of their evolutionary history, the investigation of subgenome-level evolution, including both coding and non-coding regions, has garnered considerable interest (Schnable *et al*., 2011, Schnable James *et al*., 2011, Zhao *et al*., 2017, Miao *et al*., 2020). However, despite recent advances, there remains a lack of systematically comparable studies on the evolution of non-coding regulatory regions, as well as the complex relationship between coding and non-coding regions across multi-paleopolyploidies with various and distinctive natures at the subgenome level.

We demonstrated that P. somniferum exhibited an unbiased fractionation pattern between its two subgenomes at the gene level following the recent WGD. Besides coding regions, our results also showed that the fate of the retained CNEs followed the unbiased fractionation model in *P. somniferum*, indicating the fractionation model could have the same implication for both coding and non-coding regions. To investigate this further, we included maize and soybean along with *P. somniferum* in the analysis and examined three comparable elements: distances to genes, different WGD occurrence times and different fractionation patterns at the gene level. Using the length ratios of the retained CNEs as an indicator, we conducted an intensive analysis and arrived at the following three conclusions: (1) the proportion of CNEs decreased with increasing timing of WGD events and distance of CNEs to their related genes; (2) divergent evolutionary patterns in both coding and non-coding regions between duplicates and singletons in all subgenomes; (3) fractionation pattern played a synergistic role in both coding regions and non-coding regions. Importantly, the fractionation fate of the CNEs (putative regulatory sequences) was only associated with the fate of gene loss, regardless of the time of WGD occurrence or the distance to genes. Our study focused on duplicates and singletons within diverse paleopolyploid contexts and provided insights into the evolutionary trajectory of subgenomes in coding and non-coding sequences.

### The evolvability of CNEs and the contribution of their variations to the divergence in gene expression patterns

Divergence in gene expression patterns and underlying genomic regulatory elements is known to promote adaptation and speciation, particularly among closely related species (King and Wilson, 1975, Romero *et al*., 2012, Yang and Wang, 2013, Li *et al*., 2020a). The objective of this study is to investigate the evolvability of CNEs and their contribution to the divergence in gene expression patterns. To achieve this, transcriptomic evolution was first quantitatively evaluated using PCCs to distinguish different evolutionary processes. Then, coding region variations and CNEs retained length ratios were integrated into the evolutionary framework analysis.

Our analysis showed that, at the global level of duplicates, the coding regions remained nearly identical despite the differentiation in transcriptional profiles. Conversely, variations in CNEs were consistent with transcriptional divergence and enriched specific motifs to respond to different functions of distinct transcriptional profiles. Furthermore, the divergence of gene expression patterns was faster than the divergence of gene coding sequences in duplicates (with the relative number of divergent genes as the indicator). These findings provide additional evidence for the hypothesis that regulatory variations, coordinated with transcriptomic variations, play a crucial role in governing phenotypic divergence (Wittkopp and Kalay, 2012).

Our observations and previous research have proposed intrinsically biased properties of duplicates and singletons, such as CNEs deficient genes having a decreased tendency to retain two copies after the WGD event (Schnable *et al*., 2011). A comparison of the coding and non-coding features for duplicates and singletons showed that CNEs variations were more associated with transcriptome evolutionary processes than singleton status. In contrast, relaxed selection on coding regions may be more important for singletons status (Figure 3f **and** Figure 3g, **Figure S11 and Figure S12**).

CNEs variations caused more rapid transcriptomic evolution after the WGD event than variations in gene coding sequences, indicating that speciation or adaptation might be better explained by regulatory and transcriptomic divergence. In contrast, gene losses might depend more on relaxed selection in coding regions.

### Co-regulation and co-evolution of *PSMT1-CYP719A21* cluster

The BIA gene cluster, which controls the biosynthetic pathways of morphine and noscapine, represents a notable example of transcriptional and phenotypic evolution associated with WGD events (Yang *et al*., 2021). For noscapine biosynthesis, the initial and committed step in this gene cluster is the O-methylation of (S)-scoulerine to (S)-tetrahydrocolumbamine by PSMT1, followed by its conversion to (S)-canadine by CYP719A21 (Winzer *et al*., 2012). Moreover, the *PSMT1-CYP719A21* clusters were adjacent in *P. somniferum* and *P. rhoeas*, with a specific structure of head-to-head in opposite directions (Yang *et al*., 2021). The results indicate that this structure may have been preferentially selected and maintained via co-regulation and co-evolution, as a common cis-element is shared by *PSMT1* and *CYP719A21*, which is supported by the observation that most CNEs and all ACRs are exclusively located in this specific intergenic region within the 30 kb genomic environment surrounding the *PSMT1-CYP719A21* cluster.

*PSMT1* and *CYP719A21* are closely located in *Papaver* species and the *Macleaya cordata* genome, indicating that they may be ancient seeding genes of the BIA gene cluster along with other collinearity genes (Li *et al*., 2020b, Yang *et al*., 2021). However, unlike the other putative ancient seeding genes (Yang *et al*., 2021), only a few CNEs were found flanking *PSMT1* and *CYP719A21*, which appear to be residual evolution traces. It is possible that a gene may not have any upstream CNEs due to rapid evolution and translocation. However, a gene may remain in the same genomic region if it contains many upstream CNEs. Therefore, these results provide evidence for the hypothesis that the *PSMT1-CYP719A21* cluster was among the earliest to be located in the BIA gene cluster (Li *et al*., 2020b), while the others may have undergone translocations as well.

Moreover, the “residual traces (retained CNEs)” were found to be enriched with root-specific functions, explaining the high expression levels observed in the root tissue of *P. rhoeas*. Furthermore, stem specificity in the *P. somniferum* of the *PSMT1-CYP719A21* cluster may be related to a stem-specific ACR (Figure 6b). Meanwhile, partial overlap of CNEs with this ACR suggests that the shifted expression profiles of the *PSMT1-CYP719A21* cluster might be relevant to the origin of a novel stem-specific ACR, using previous cis-elements as templates. These findings demonstrate that variations in CNEs can result in transcriptional and phenotypic evolution and provide a systematic and credible example of genomic, transcriptomic, and epigenetic data to explain the co-evolution of the BIA gene cluster.

### Future perspectives

In this study, we have highlighted the significance of CNEs variations and transcriptomic evolution in plant science. Our findings have revealed the evolutionary trajectories of these features, including variations in the coding regions, subsequent to WGD events. Nonetheless, our study has two limitations that warrant further investigation.: (1) While CNEs could be treated as proxies of cis-elements and residual traces of evolution, the lack of actual epigenomic data integration hampers further research. Our study was constrained by the limited availability of ATAC-seq data for *P. somniferum* and the absence of any epigenomic data for *P. rhoeas*. (2) Investigating the underlying phenotypic changes resulting from non-coding region variations requires more systematic analysis using population-level data of *P. somniferum*. *Papaver* species have undergone unique adaptation and evolution, and could be considered a promising new model system for addressing a variety of evolutionary questions.

## Materials and Methods

### Data collection

Genome sequences, annotation files and expression level data of *P. setigerum*, *P. somniferum* and *P. rhoeas* were obtained from (https://github.com/xjtu-omics/Papaver-Genomics) (Yang *et al*., 2021). ATAC-seq data and another set of expression level data from *P. somniferum* was obtained from (Jia *et al*., 2023). Expression level data from six tissues (fine root, capsule, leaf, petal, stamen, and stem) of *P. setigerum* and *P. rhoeas*. Expression level data from fifteen samples of *P. somniferum*. Maize and soybean subgenome sequences were obtained from (Brohammer *et al*., 2018) and (Zhao *et al*., 2017), respectively. Genome sequences and annotation files of maize (NCBIv4.34), sorghum (NCBIv3.42), soybean (v2.1.51), and common bean (v1.0.51) were obtained from Ensembl Plants (Howe *et al*., 2020).

### Identification of two subgenomes of *P. somniferum*

524,217 and 403,705 gene pairs paralogous in *P. somniferum* and *P. rhoeas*, and 434,700 gene pairs homologous between *P. somniferum* and *P. rhoeas*, were identified using BLASTP v2.12.0 with parameters-evalue 1e-5 and -max_target_seqs 11 (Altschul *et al*., 1997). BLASTP hits with scores < 100 were discarded. Tandemly duplicated genes were identified with MCScanX (Wang *et al*., 2012) with default parameters, using the 405,770 and 312,928 remaining BLASTP hits of *P. somniferum* and *P. rhoeas*, respectively, as input. Gene pairs which contained a tandem duplicate were discarded. Next, we used WGDI v0.4.7 (Sun *et al*., 2021) with default parameters to create a set of 784 interspecies syntenic blocks and 521 intraspecies syntenic blocks of *P. somniferum* and to calculate the evolutionary distance between them (as Ka and Ks values). We retained only those intraspecies syntenic blocks containing > 10 gene pairs and Ks values lower than 0.2 (the indicator of this recent WGD event) (Guo *et al*., 2018).

Finally, *P. somniferum* subgenomes were reconstructed on the basis of differences between subgenome-specific k-mers (Jia *et al*., 2022) (method 1) and on differences in the retention rate of genes in each block (Zhao *et al*., 2017) (method 2). For method 1, we identified subgenome-specific k-mers using SubPhaser v1.2.5 (Jia *et al*., 2022) with default parameters, setting chr1 and chr6, and chr9 and chr11 as homologous chromosome sets. By default, SubPhaser partitions each chromosome into 1 Mb bins and then classifies each bin into one of two categories (‘SG1’ or ‘SG2’) on the basis of differential k-mer use (p-value < 0.05, Fisher’s exact test) (**Figure S1, Table S1**). By combining these classifications with the set of interspecies syntenic blocks, we could reconstruct two subgenomes, one comprising 239 syntenic blocks classified as SG1 and one comprising 197 syntenic blocks classified as SG2. The final results were manually checked and corrected. For method 2, the ratio of *P. somniferum* genes to *P. somniferum* genes with *P. rhoeas* synteny in each duplicated block was calculated. Blocks were then assigned to one of two subgenomes on the basis of whether they were more or less fractionated (higher or lower proportion of *P. somniferum* genes to *P. somniferum* genes with *P. rhoeas* synteny, respectively).

### Identification of conserved non-coding elements

CNEs within the maize, soybean, *P. setigerum* and *P. somniferum* genomes were identified by pairwise comparison of maize with sorghum, soybean with common bean, *P. setigerum* with *P. rhoeas*, and *P. somniferum* with *P. rhoeas*, in each case using the CNS Discovery Pipeline v.3.0 (Turco *et al*., 2013) with subgenome scaffolding. All genomes were repeat-masked prior to input, using RepeatModeler v2.0.2 (Flynn Jullien *et al*., 2020) and RepeatMasker v4.1.2 (Smit *et al*., 2015) with default parameters.

To complement this approach, CNEs between the *P. somniferum* genome and *P. rhoeas* genome were also identified using dCNS v0.4 (Song *et al*., 2021) with the following parameters (the genome masked parameter “-f 26 for *P. somniferum*, 35 for *P. rhoeas*”, k-mer counting parameter “-m 25”, and CNEss discover parameter “-A 2 -w 45 -c 50 - s 10 -k 0.00501 −l 0.34725”).

### Gene expression level analyses

Gene expression data of both *P. somniferum* and *P. rhoeas* were normalised using the R/Bioconductor package preprocessCore v1.48.0 (Bolstad *et al*., 2003) and then log_2_-trasformed. For each gene pair, we required that both genes were detectably expressed in at least one tissue (maximum normalised expression >= 1), prior to log_2_-trasformation.

To compare the expression profiles of genes and their orthologues, we first established a set of 2,909 single-copy orthologues common to the three *Papaver* species using Orthofinder v2.2.6 with parameters-S diamond-M msa (Emms and Kelly, 2015). We then calculated Pearson correlation coefficients for the pairwise comparisons of *P. somniferum* with *P. rhoeas*. The mean values of PCCs were used as the cutoff values for R_O1,O2_. Then, the PCCs between duplicates and their ancestral genes and between singletons and their ancestral genes were calculated. Finally, the evolutionary processes of duplicates and singletons were classified by each PCC value with R_O1,O2_, as previously described.

For the cluster tree of CNEs retained lengths of different evolutionary processes, the distributions of CNEs retained lengths were cut using 1 as unit. For the cluster tree of Ka/Ks values, the distributions of Ka/Ks values were cut using 0.01 (range 0-0.6). The values were normalised to the same range by dividing the count of each box by the total count.

### ATAC-seq data analysis and motif analysis

The number of CNEs bases within an accessible chromatin region (ACR) was calculated as follows. First, fifteen bam files of five tissues (fine root, tap root, leaf, petal, and stem) from a previous study of *P. somniferum* (Jia *et al*., 2023) with those from the same tissues merged. Peaks were called with MACS2 v2.2.6 with parameters-f BAMPE -q 0.05 -g 2408287349 --keep-dup all (Zhang *et al*., 2008). Second, ACRs within 1 kb regions upstream of TSS for a gene on each of the two subgenomes were extracted using BEDtools intersect v2.30.0 (Quinlan and Hall, 2010). The number of ACR bases overlapping each CNEs were also calculated using BEDtools intersect. The ‘multiBamSummar’ script from deepTools v2.0 (Ramírez *et al*., 2014) was used to count the number of ATAC-seq reads of CNEs and Non_CNEs regions within 1 kb regions upstream of TSS and intronic regions across five tissues. The values were then normalised with length (sequence lengths greater than 30 were used for analysis).

Motifs enrichment was calculated as follows. First, coordinates of upstream 1 kb regions for both_conservation (both duplicates are conserved with their ancestral genes) and both_specialization (both duplicates are divergent with their ancestral genes)-related genes were extracted. Second, BEDtools was used to obtain a list of CNEs within the above regions. Third, BEDtools was used to randomly shuffle CNEs annotations within 1 kb regions for the background of each evolutionary process (Quinlan and Hall, 2010). Fourth, the ‘annotatePeaks.pl’ function of the HOMER package v4.11 (Heinz *et al*., 2010) was used for motifs annotations on each CNEs lists or random CNEs lists. The motifs files were obtained from the previous study (O’Malley *et al*., 2016). Fifth, enrichment motifs were selected based on two principles: (1) the number of real observed annotations located at least above 99% of random distribution; and (2) the number of real observed annotations at least 1.25-fold of the median of random distribution. Sixth, each motif was annotated using R package UniprotR (v2.2.0) (Soudy *et al*., 2020).

### Enrichment analyses

KEGG enrichment analysis was conducted using TBtools v1.098761 (Chen *et al*., 2020). GO enrichment analysis was conducted using UniprotR v2.2.0 (Soudy *et al*., 2020) with a adjusted p-value threshold of 0.01.

### Visualization

Unless otherwise specified, all figures were conducted in R v3.6.2 (Team, 2013), with packages pheatmap v1.0.12 (Kolde, 2019), VennDiagram v1.7.1, ggplot2 v3.3.2 (Wickham, 2016), and ChIPseeker v1.22.1 (Yu *et al*., 2015). Genome browser images were produced using Integrative Genomics Viewer (IGV) v2.8.10 (Thorvaldsdóttir *et al*., 2013).

## Acknowledgements

We thank Yubin Yan and Panpan Lei for the insightful suggestions and Tun Xu for the advice on manuscript preparation and data analysis. This work was supported by the National Natural Science Foundation of China (32125009, 32070663, 32200510) and the Key Construction Program of the National ‘985’ Project.

## Code availability

All scripts used for these analyses are available at https://github.com/StuYuXu/poppy_subgnome.

## Short legends for supporting information

**Figure S1.** Chromosomal characteristics of opium poppy. From outer to inner circles (1–7): (1) Rough subgenome assignments for each chromosome. (2) Significant enrichment of subgenome-specific k-mers. Yellow bars indicate significant enrichment of subgenome1-specific k-mers. Blue bars indicate significant enrichment of subgenome2-specific k-mers. (3) Relative subgenome-specific k-mers for each bar or bin. (4) Count for subgenome1-specific k-mer set. (5) Count for subgenome2-specific k-mer set. (6) Density of long terminal repeat retrotransposons (LTR-RTs), which were enriched to those subgenome-specific k-mers. Gray indicates non-specific LTR-RTs. (7) Homoeologous blocks were identified in homoeologous chromosome sets. These homoeologous chromosome sets were used as configuration to identify subgenome-specific k-mers. This figure was the output of subphaser.

**Figure S2.** Genes retained ratio of each *P. rhoeas* chromosome using the k-mer differences method. The percentage (Y-axis) indicates the number of genes retained for subgenome1 (blue), subgenome2 (red) and both subgenomes (green) based on 100 genes sliding windows along each *P. rhoeas* chromosome (X-axis).

**Figure S3.** Genes retained ratio of each *P. rhoeas* chromosome using the retained rates differences method. The percentage (Y-axis) indicates the number of genes retained for subgenome1 (blue), subgenome2 (red) and both subgenomes (green) based on 100 genes sliding windows along each *P. rhoeas* chromosome (X-axis).

**Figure S4.** Comparisons of the overall gene expression level (a) and expression breadths (b) between duplicates and singletons in subgenome1 and subgenome2 reconstructed under two methods in seven different tissues. The total gene expression level was calculated via log_2_ (sum expression values of seven different tissues + 1). The cutoff of expression breadth was TPM >= 1, which was expressed in that tissue. The statistical analysis was conducted via the Wilcoxon test. The asterisk ** indicates adjusted p-value < 0.05, which was considered as significant under the Benjamini & Hochberg correction (FDR, false discovery rate).

**Figure S5.** Comparison of normalised overlapped ATAC-seq reads between CNEs and Non_CNEs within intronic regions. Sequences longer than 30 bp were considered for analysis. The statistical analysis was conducted via the Wilcoxon test. Asterisk ** indicates adjusted p-value < 0.05 under the Benjamini & Hochberg correction. The outlier dots were masked by ggplot2.

**Figure S6.** Comparisons of the percentage of ACRs related bases located in regions associated with the CNEs and Non_CNEs within 1 kb upstream of TSS. The leftmost bar plot indicates the relative lengths of CNEs and Non_CNEs within 1 kb upstream of TSS.

**Figure S7.** Comparison of CNEs retained length ratios between subgenome1 and subgenome2 for each chromosome. The statistical analysis was conducted via the Wilcoxon test.

**Figure S8.** Comparison of CNEs retained numbers between subgenome1 and subgenome2 for 10 different regions. The statistical analysis was conducted via the Wilcoxon test.

**Figure S9.** Comparisons of the (**a**) Ka/Ks values, (**b**) identity values, and (**c**) CNEs retained length ratios (1 kb) for different evolutionary processes-related genes in duplicates. Subgenome1 genes of Specialization_1 process and Subgenome2 genes of Specialization_2 process were merged into Specialization. Subgenome2 genes of Specialization_1 process and Subgenome1 genes of Specialization_2 process were merged into Specialization_retain. The identity value was calculated using BLAST. Statistical analysis was conducted via the Wilcoxon test or Wilcoxon paired test. “ns” indicates not significant (p-value > 0.05). The asterisk ** indicates adjusted p-value < 0.05, and * indicates p-value < 0.05. P-value < 0.05 was considered as significant, while adjusted p-value < 0.05 was considered as significant under the Benjamini & Hochberg correction.

**Figure S10.** Comparisons of the (**a**) Ka/Ks values and (**b**) CNEs retained length ratios (1 kb) for different evolutionary processes-related genes in singletons. The statistical analysis was conducted via the Wilcoxon test. Asterisk ** indicates adjusted p-value < 0.05 under the Benjamini & Hochberg correction.

**Figure S11.** The cluster tree topology of the (**a**) Ka/Ks values and (**b**) CNEs retained lengths (1 kb) for different evolutionary processes between duplicates and singletons with the median value of PCCs as the threshold. Statistical analysis was conducted via the Wilcoxon test. “ns” indicates not significant (p-value > 0.05). Asterisk ** indicates adjusted p-value < 0.05 and * indicates p-value < 0.05. P-value < 0.05 was considered as significant, while adjusted p-value < 0.05 was considered as significant under the Benjamini & Hochberg correction. The outlier dots were masked by ggplot2.

**Figure S12.** The cluster tree topology of the (**a**) Ka/Ks values and (**b**) CNEs retained lengths (1 kb) for different evolutionary processes between duplicates and singletons with the mean value of SCCs as the threshold. Statistical analysis was conducted via the Wilcoxon test. “ns” indicates not significant (p-value > 0.05). Asterisk ** indicates adjusted p-value < 0.05 and * indicates p-value < 0.05. P-value < 0.05 was considered as significant, and adjusted p-value < 0.05 was considered as significant under the Benjamini & Hochberg correction. The outlier dots were masked by ggplot2.

**Figure S13.** Comparison of normalised ATAC-seq reads contained between conserved genes and specialized genes. ATAC-seq reads were normalised by the number of reads in each gene divided by the total number of reads of all genes and multiplied by 10^6 for each ATAC-seq sample. The length was not normalised because each gene is identical (1 kb). Statistical analysis was conducted via the paired and single-sided t-test. The outlier dots were masked by ggplot2.

**Figure S14.** An example for *Pso06G01300.0* of CNEs variations associated with the origin of novel petal-specific ACR. The brown transparent boxes represent ACRs. The pink and blue transparent boxes represent CNEs regions for subgenome1 and subgenome2, respectively. The red transparent boxes represent the core CNEs-region. The orange transparent boxes represent the core CNEs-region variations across four orthologs in *P. setigerum*.

**Figure S15.** An example of CNEs positional variations associated with gene expression divergence in *P. somniferum*. The brown transparent boxes represent ACRs. The pink and blue transparent boxes represent CNEs regions for subgenome1 and subgenome2, respectively. The red transparent boxes represent the core CNEs-region.

**Figure S16.** An example of CNEs positional variations associated with gene expression divergence in *P. setigerum*. The red transparent boxes represent the core CNEs-region. The orange transparent boxes represent the core CNEs-region variations across four orthologs in *P. setigerum*.

**Figure S17.** Functional enrichment analysis (biological process) of specific enriched binding motifs for (**a**) both_conservation (**b**) both_specialization related CNEs.

**Figure S18.** Comparisons of observed values with background distribution for *HHO3*.

**Figure S19.** The genomic environment surrounding *PSMT1-CYP719A21* cluster within 30 kb.

**Table S1**. Subgenome-specific enrichments within each genome bin.

**Table S2.** Detail features of two subgenomes for *P. somniferum*.

**Table S3.** The statistics of singletons and duplicates of each subgenome.

**Table S4**. Inconsistent gene lists of group assignments based on two different methods.

**Table S5.** Complete list of CNEs identified between *P. somniferum* and *P. rhoeas* using the CNS Discovery Pipeline v.3.0.

**Table S6.** The quantile-based length distribution of CNEs for three different species.

**Table S7.** Complete list of CNEs identified between *P. somniferum* and *P. rhoeas* using dCNS software.

**Table S8.** Enrichment of regulatory sequences with CNEs.

**Table S9.** Details of evolutionary trajectories for *P. somniferum* duplicates with the mean value of PCCs as the threshold.

**Table S10.** Details of evolutionary trajectories for *P. somniferum* singletons with the mean value of PCCs as the threshold.

**Table S11.** Details of evolutionary trajectories for *P. somniferum* duplicates and singletons with the mean value of selection pressures as the threshold.

**Table S12.** KEGG enrichment results of four types of evolutionary trajectories for duplicates.

**Table S13.** KEGG enrichment results of four types of evolutionary trajectories for singletons.

**Table S14.** Functional annotations of specific enriched binding motifs for both_conservation related CNEs.

**Table S15.** Functional annotations of specific enriched binding motifs for both_specialization related CNEs.

**Table S16.** CNEs lists for *PSSDR1*, *CYP719A21*, *PSMT1* and *SALR*.

