## Supplementary figures and images for "Evolutionary analysis of conserved non-coding elements subsequent to whole-genome duplication in opium poppy"

### Figure S1.png

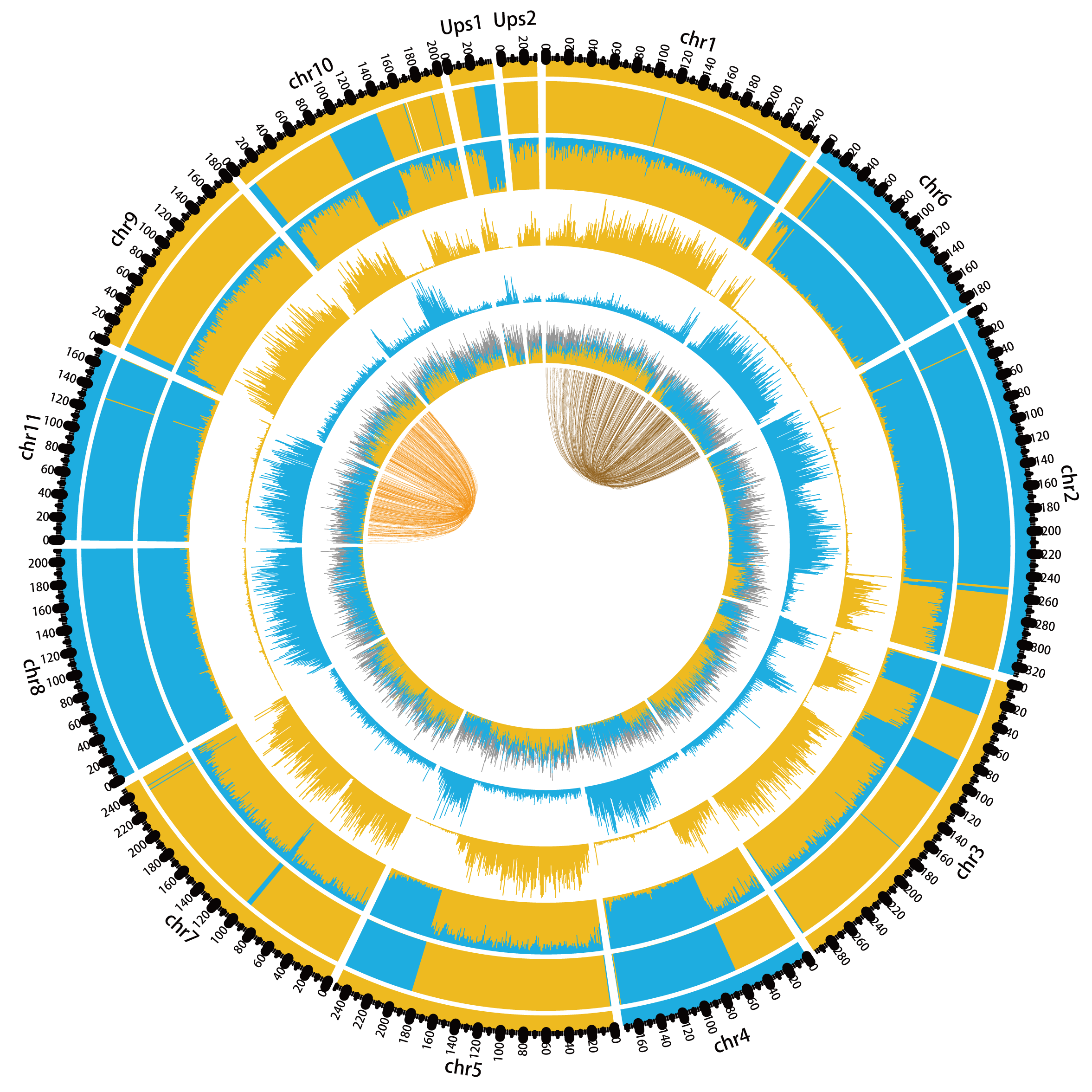

### Figure S2.tiff

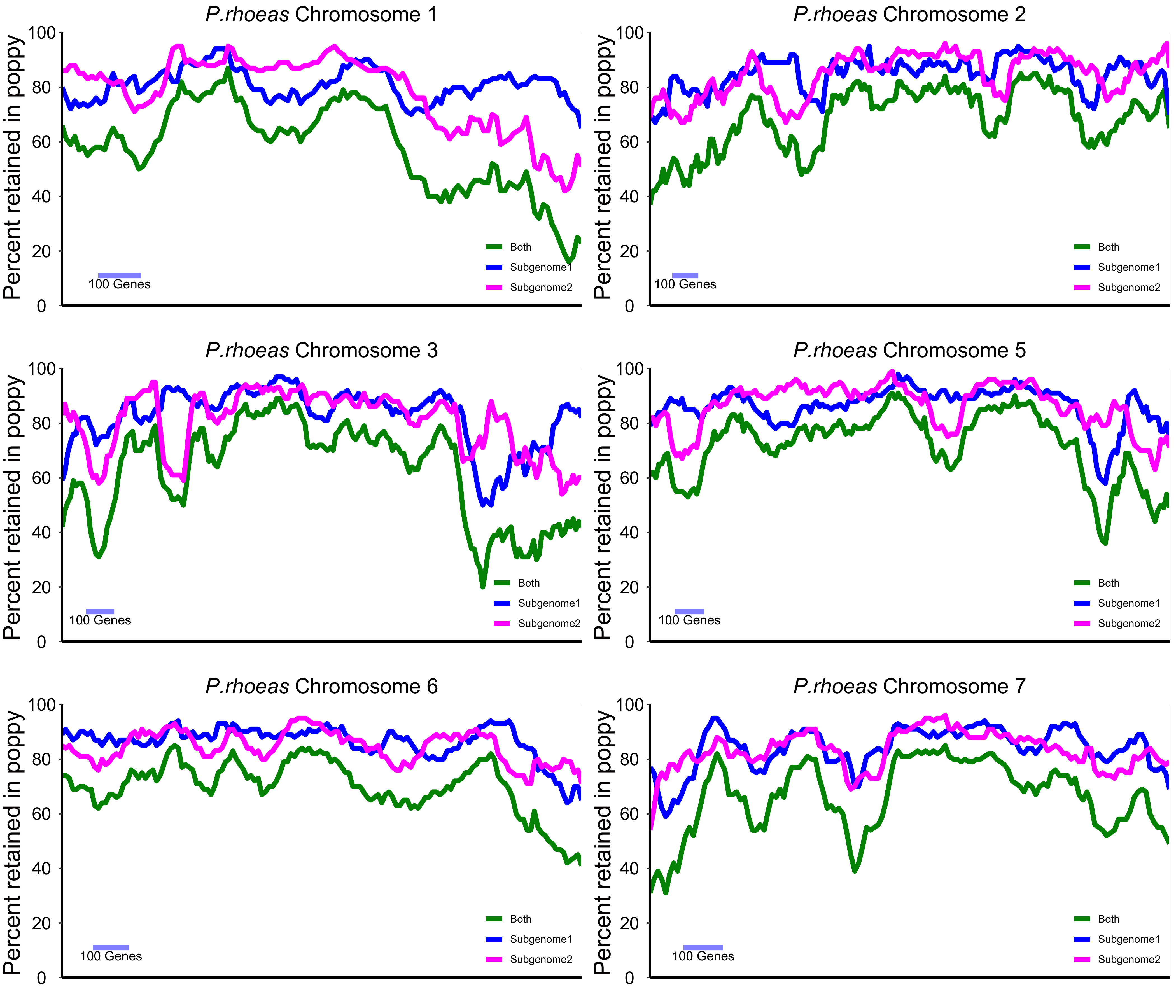

### Figure S3.tiff

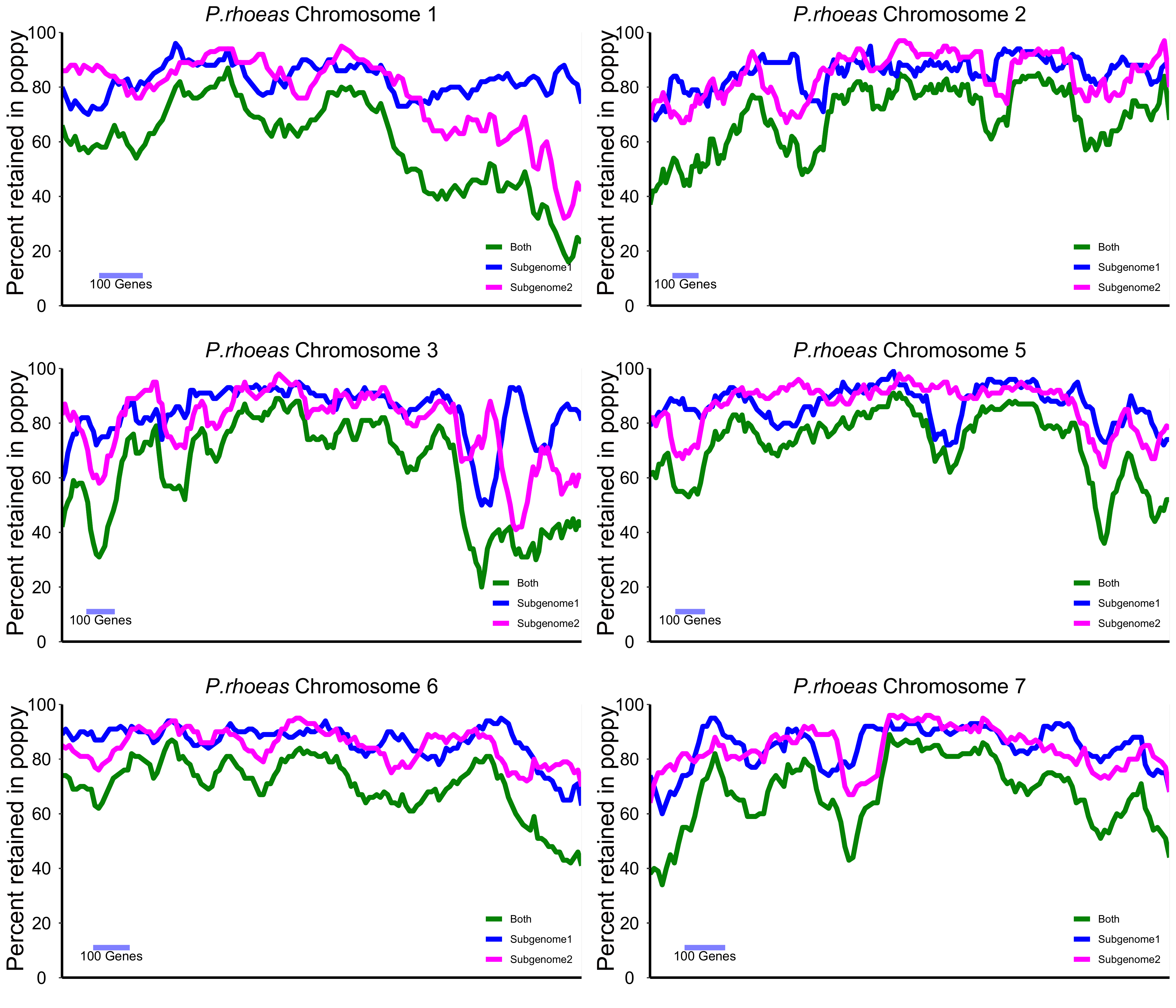

### Figure S4.tiff

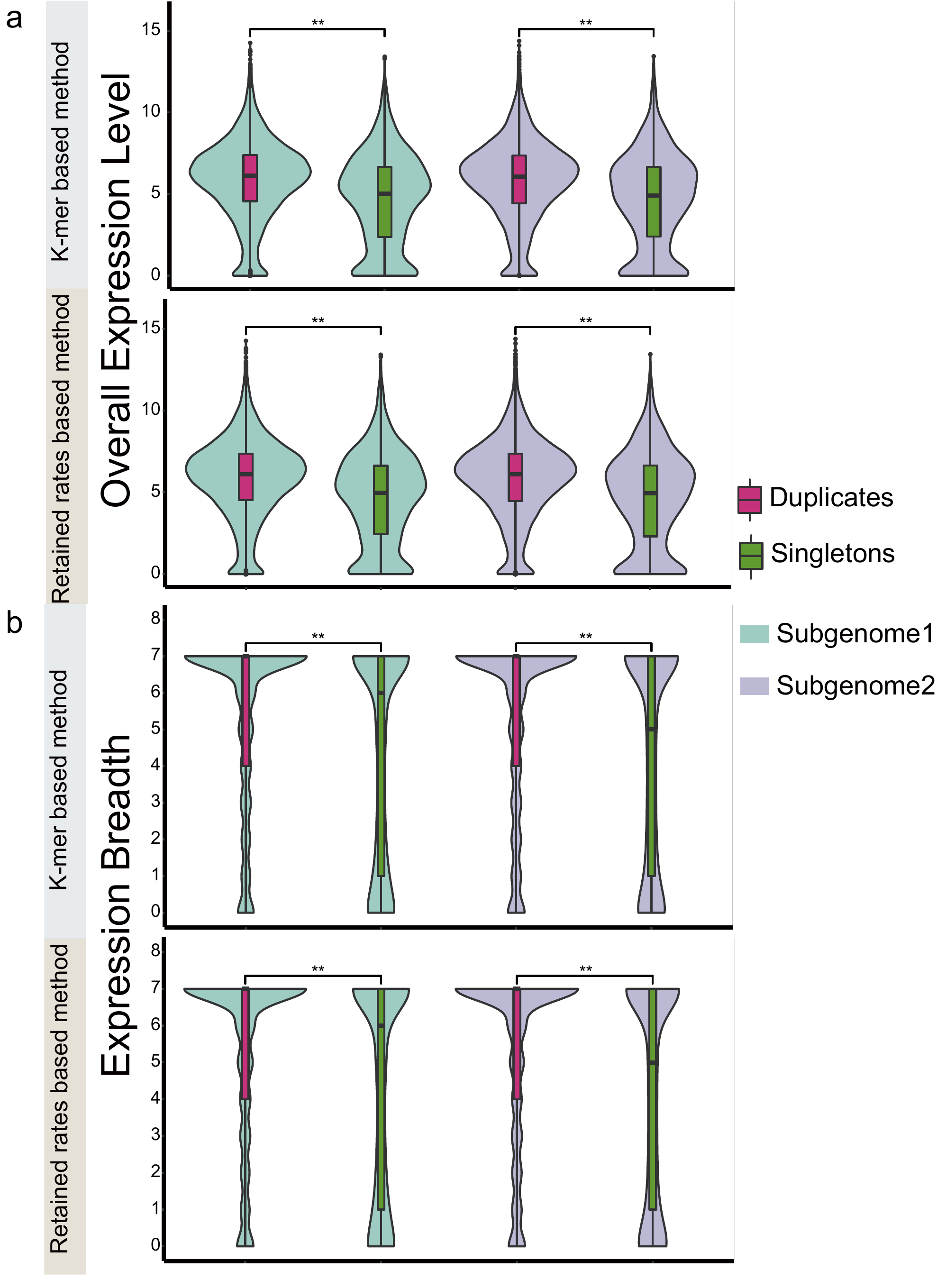

### Figure S5.tiff

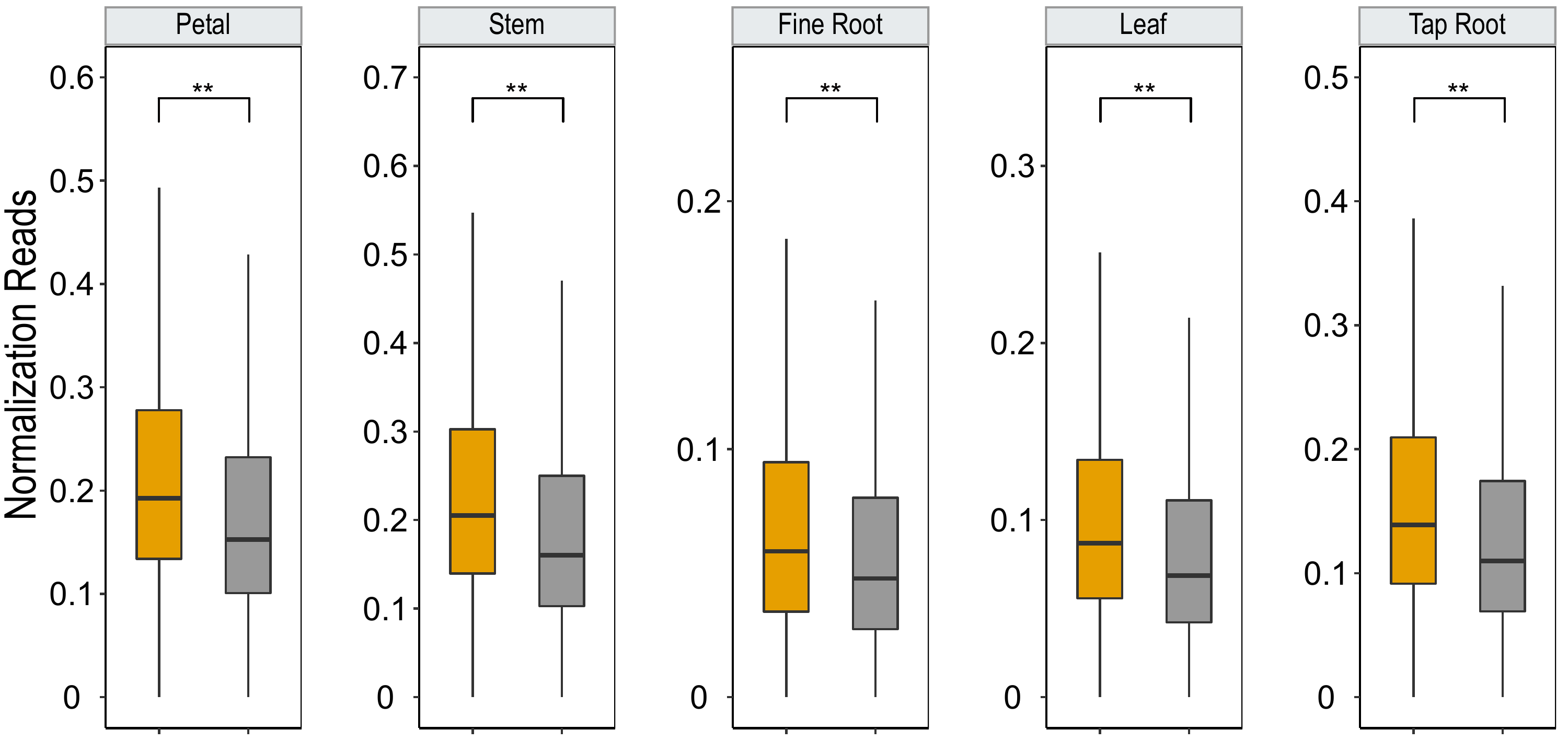

### Figure S6.tiff

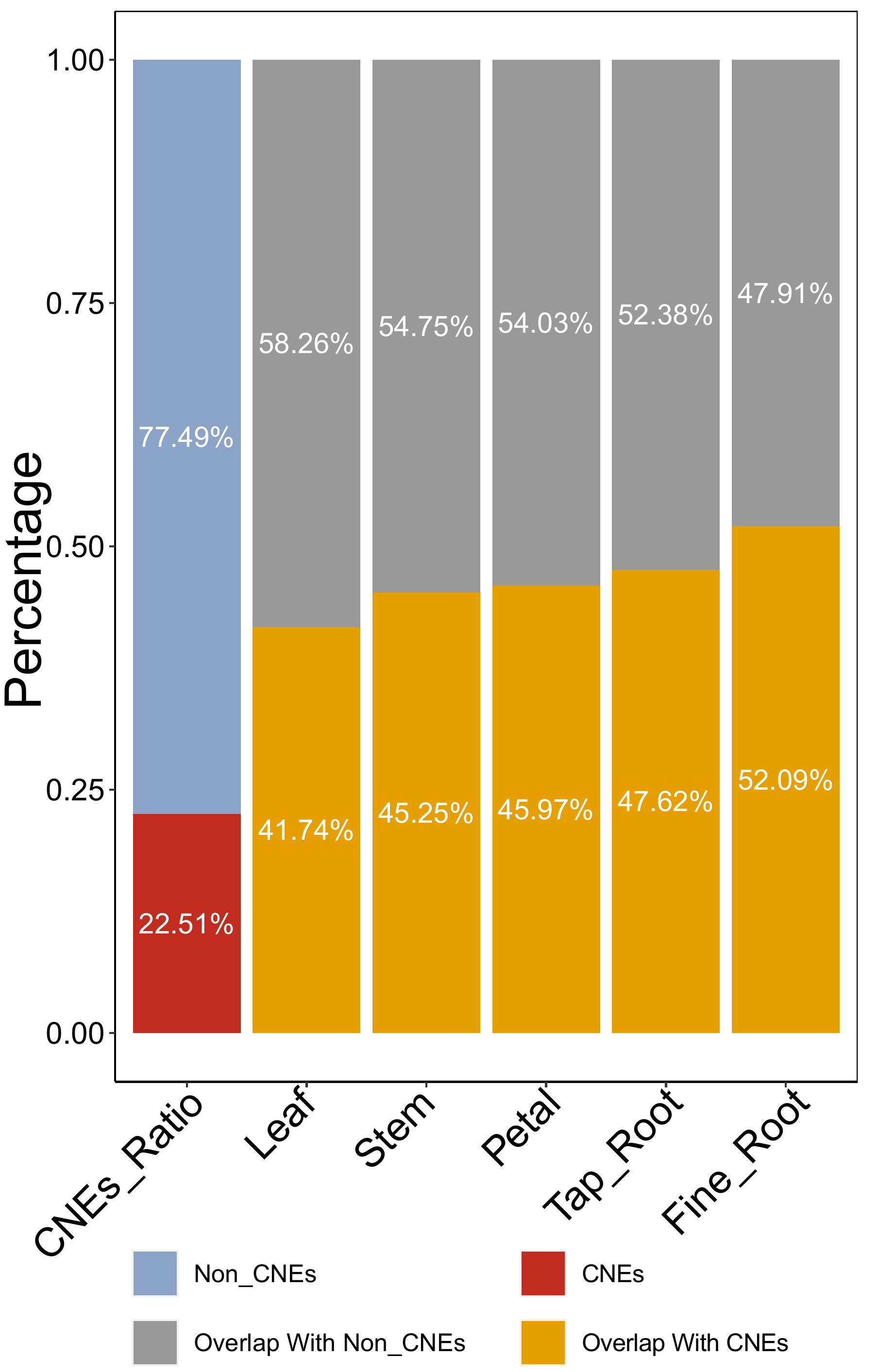

### Figure S8.tiff

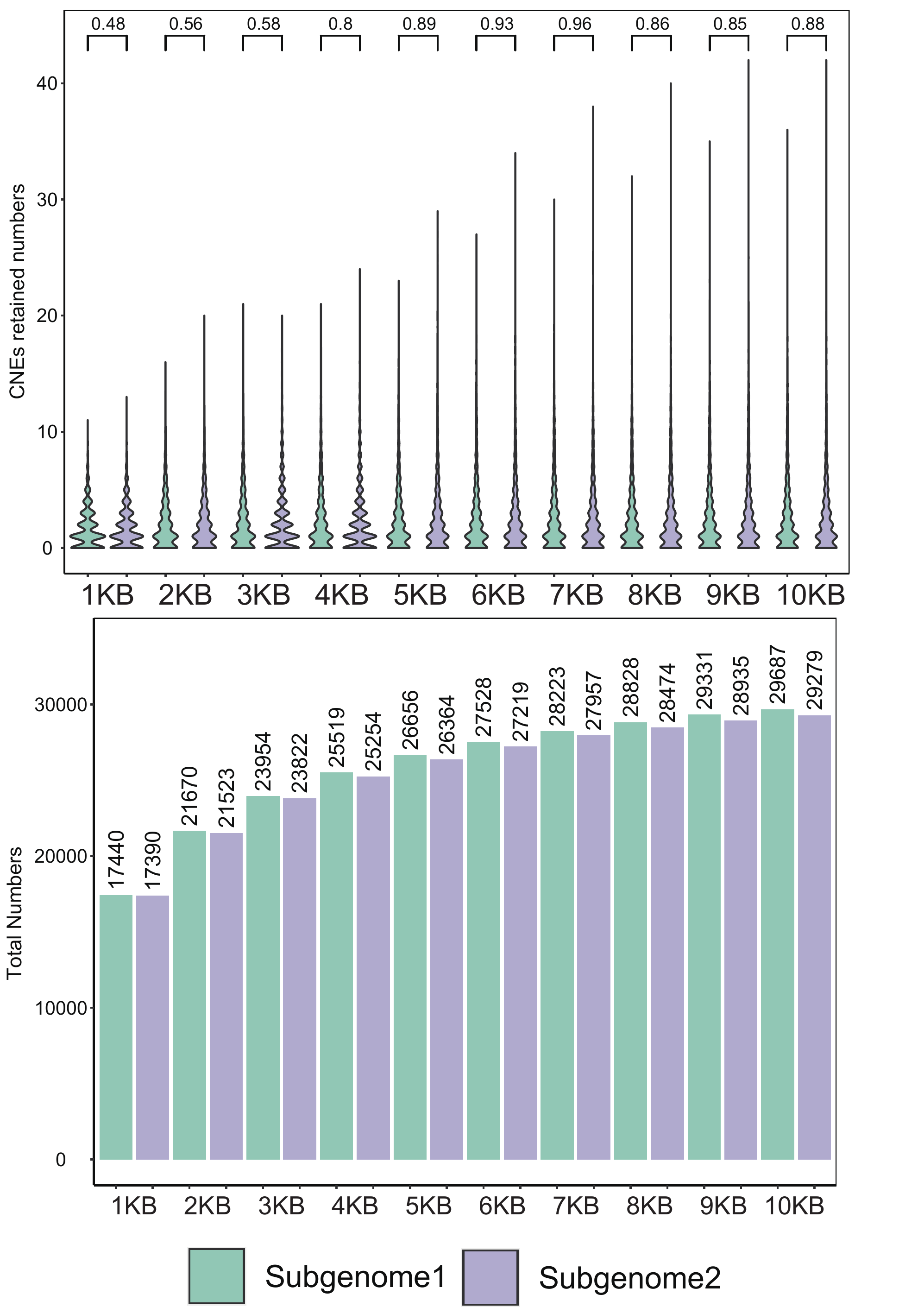

### Figure S9.tiff

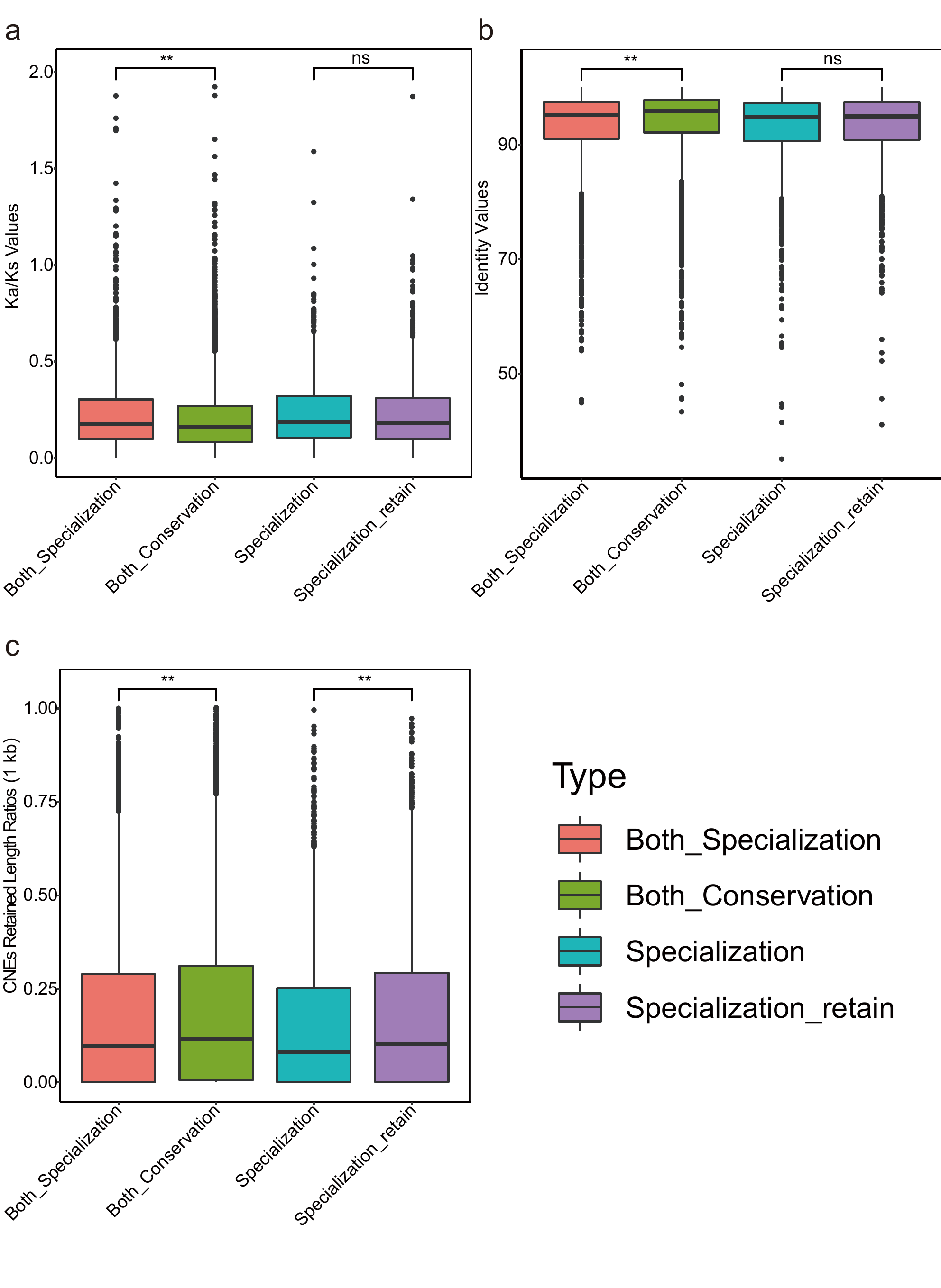

### Figure S10.tiff

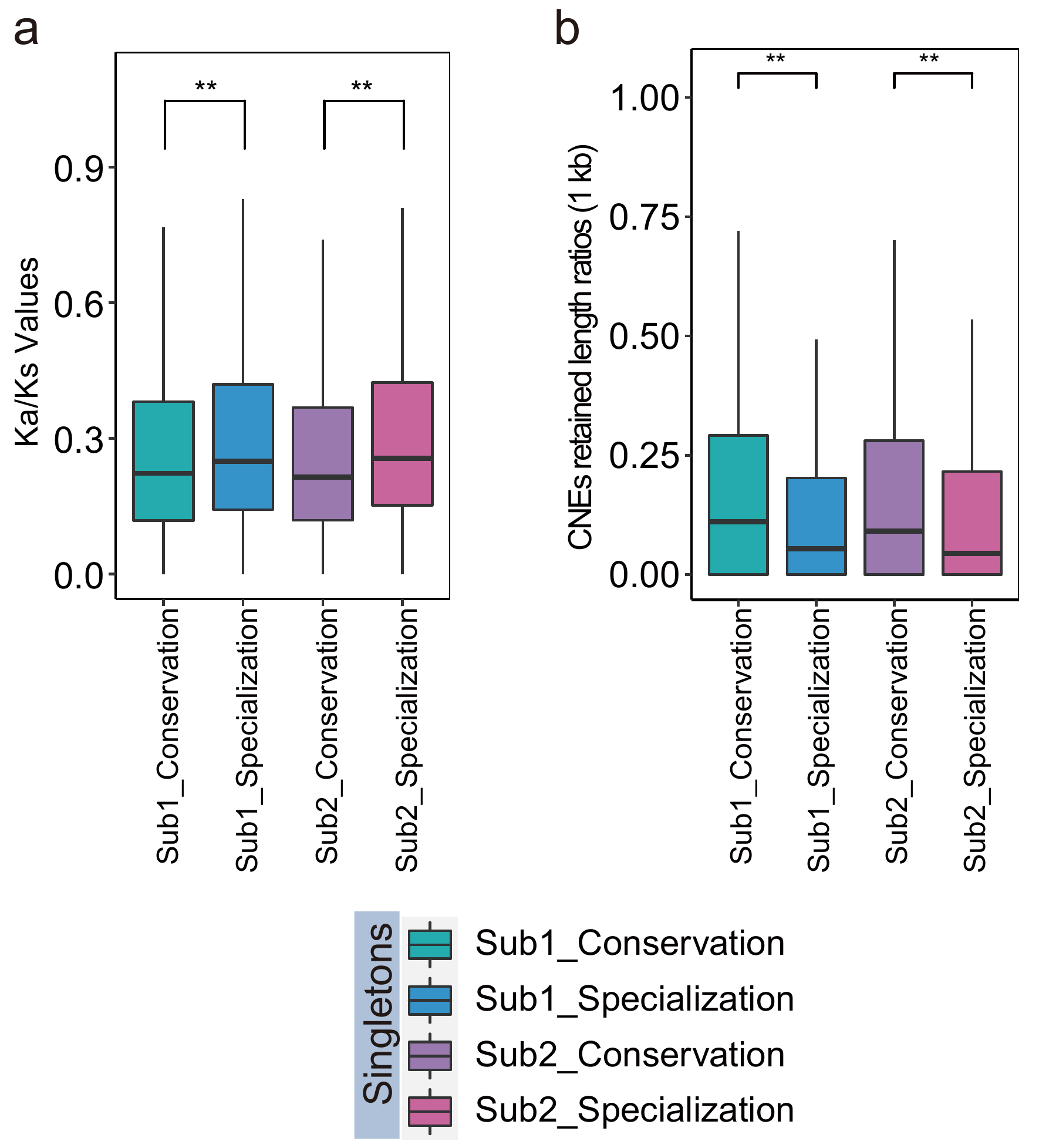

### Figure S11.tiff

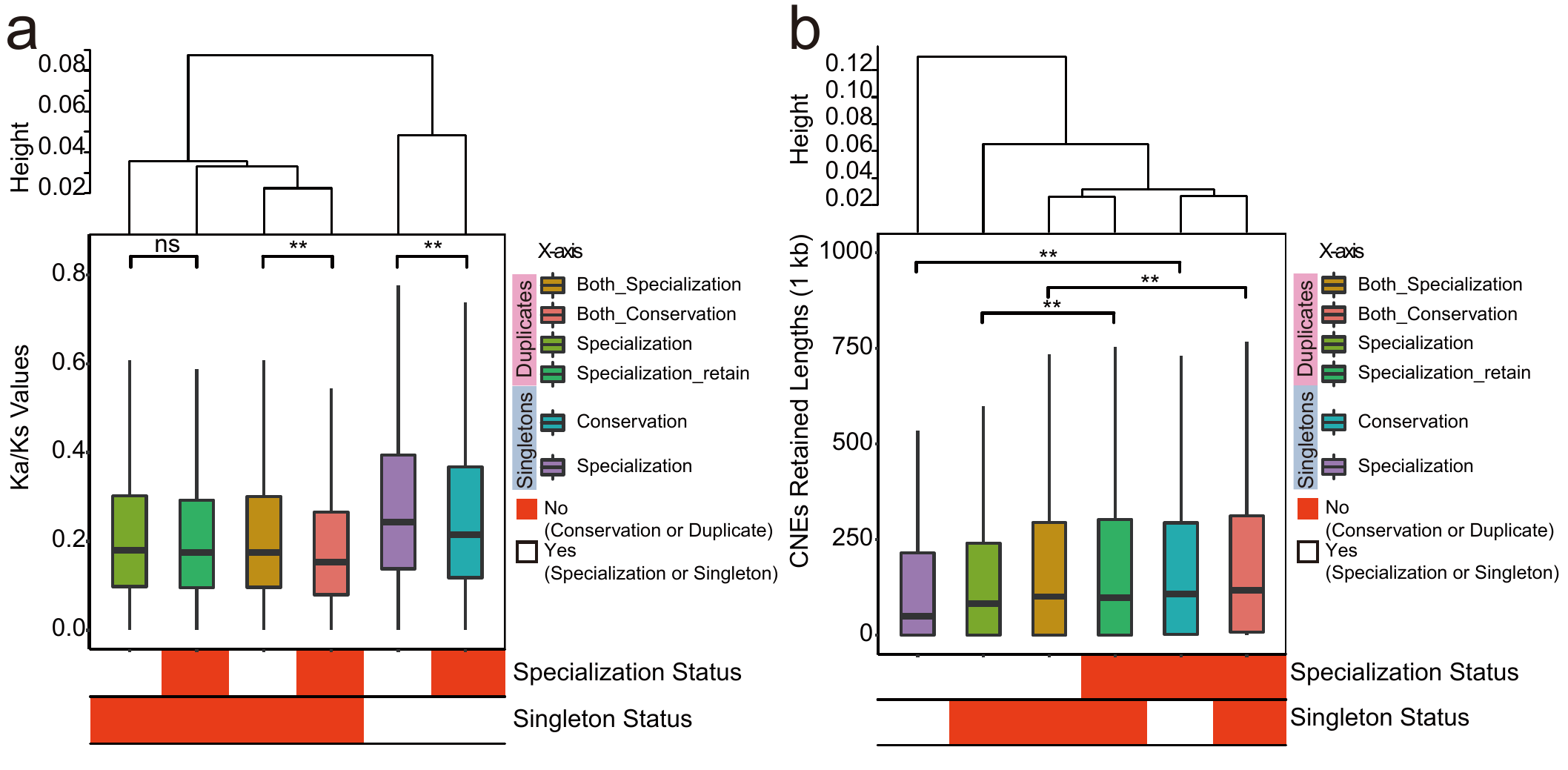

### Figure S12.tiff

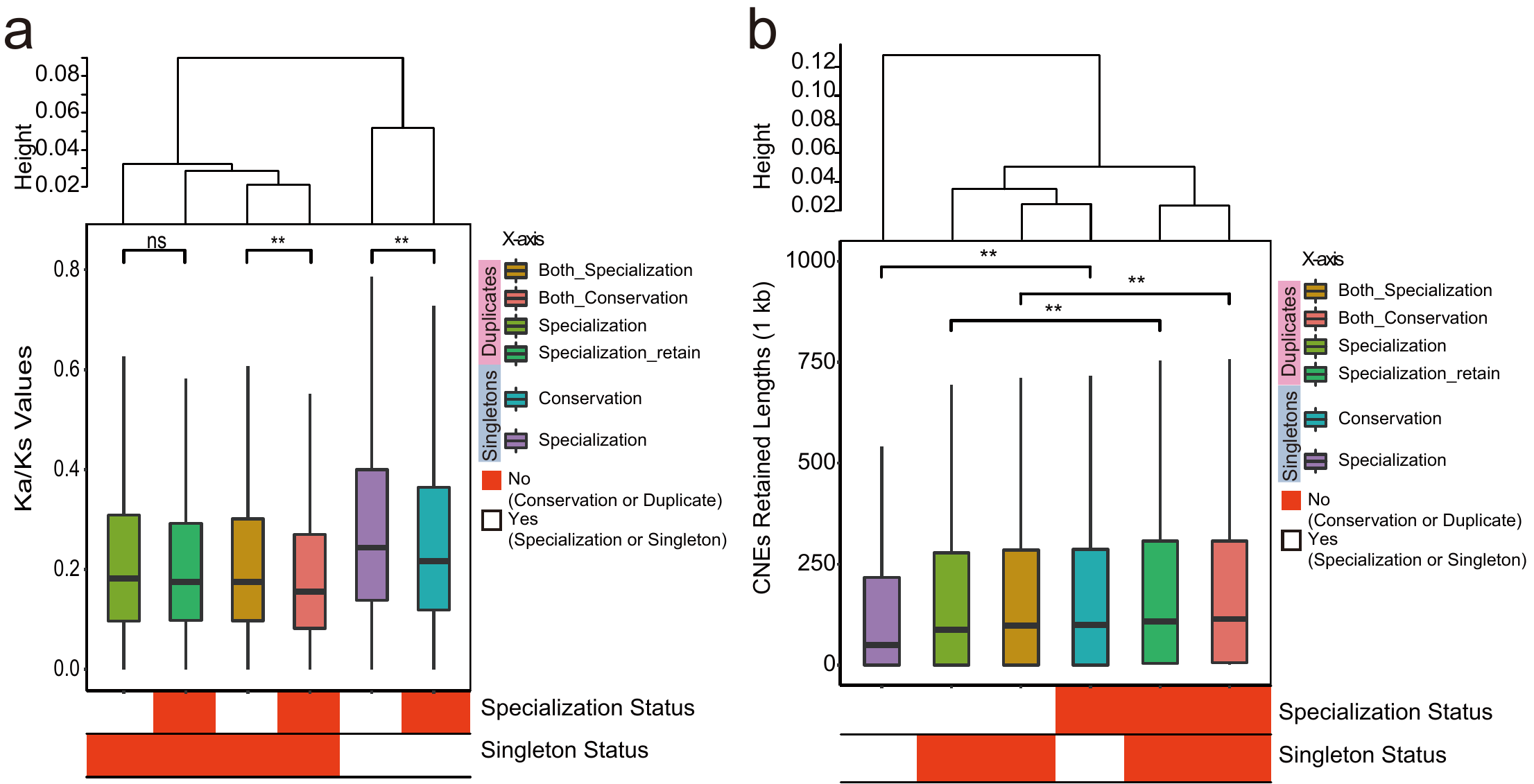

### Figure S13.tiff

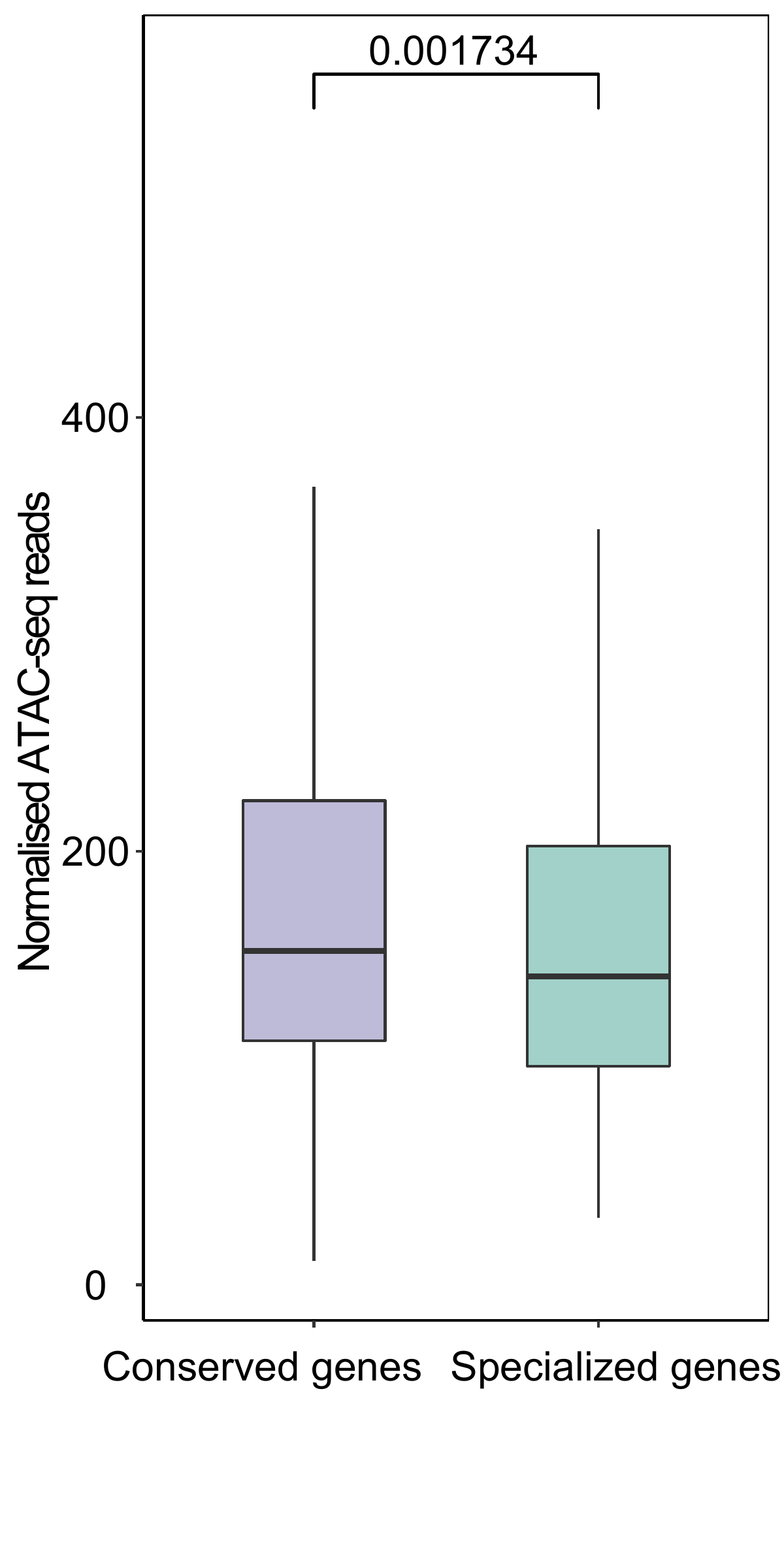

### Figure S14.tiff

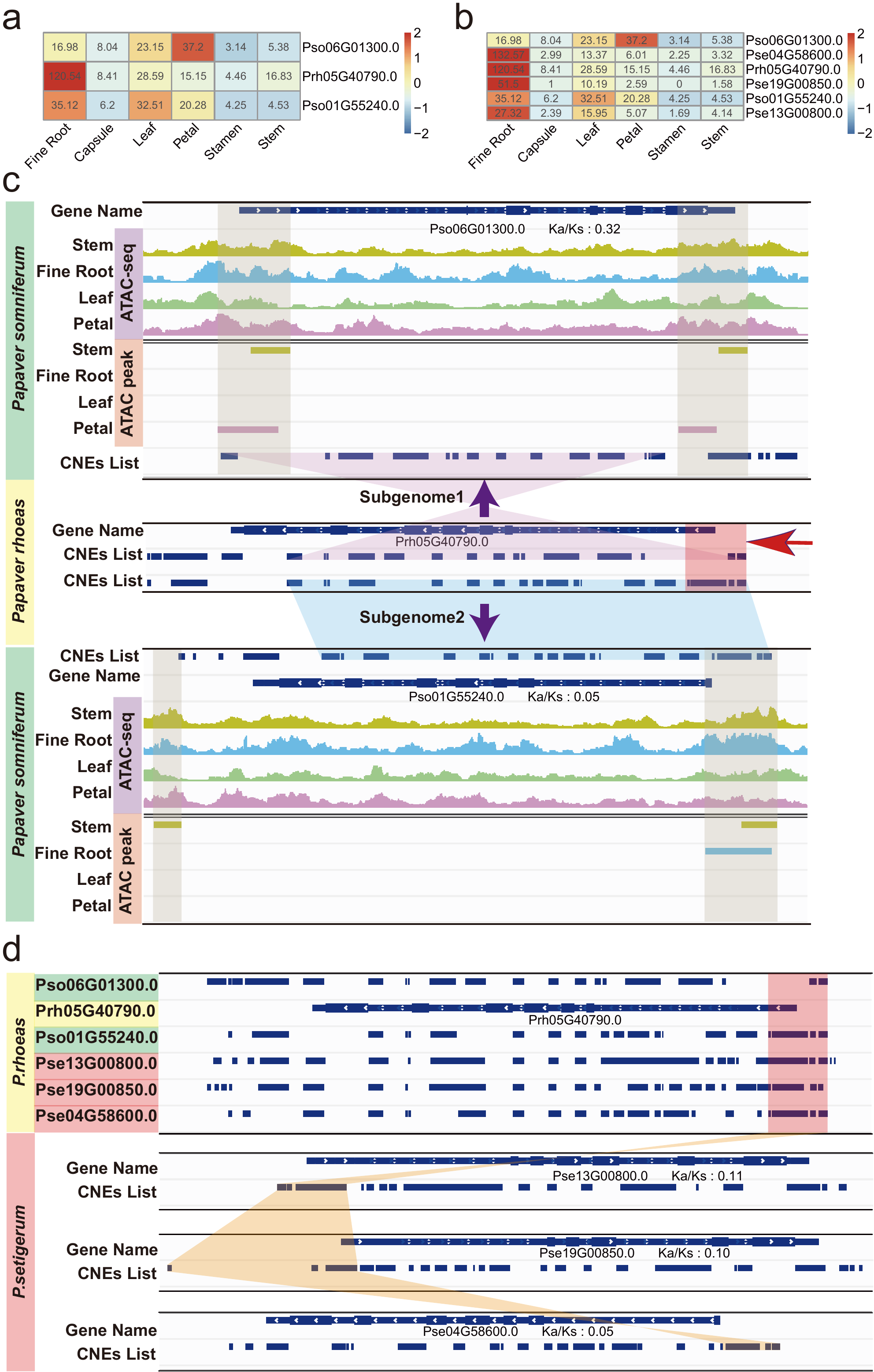

### Figure S15.tiff

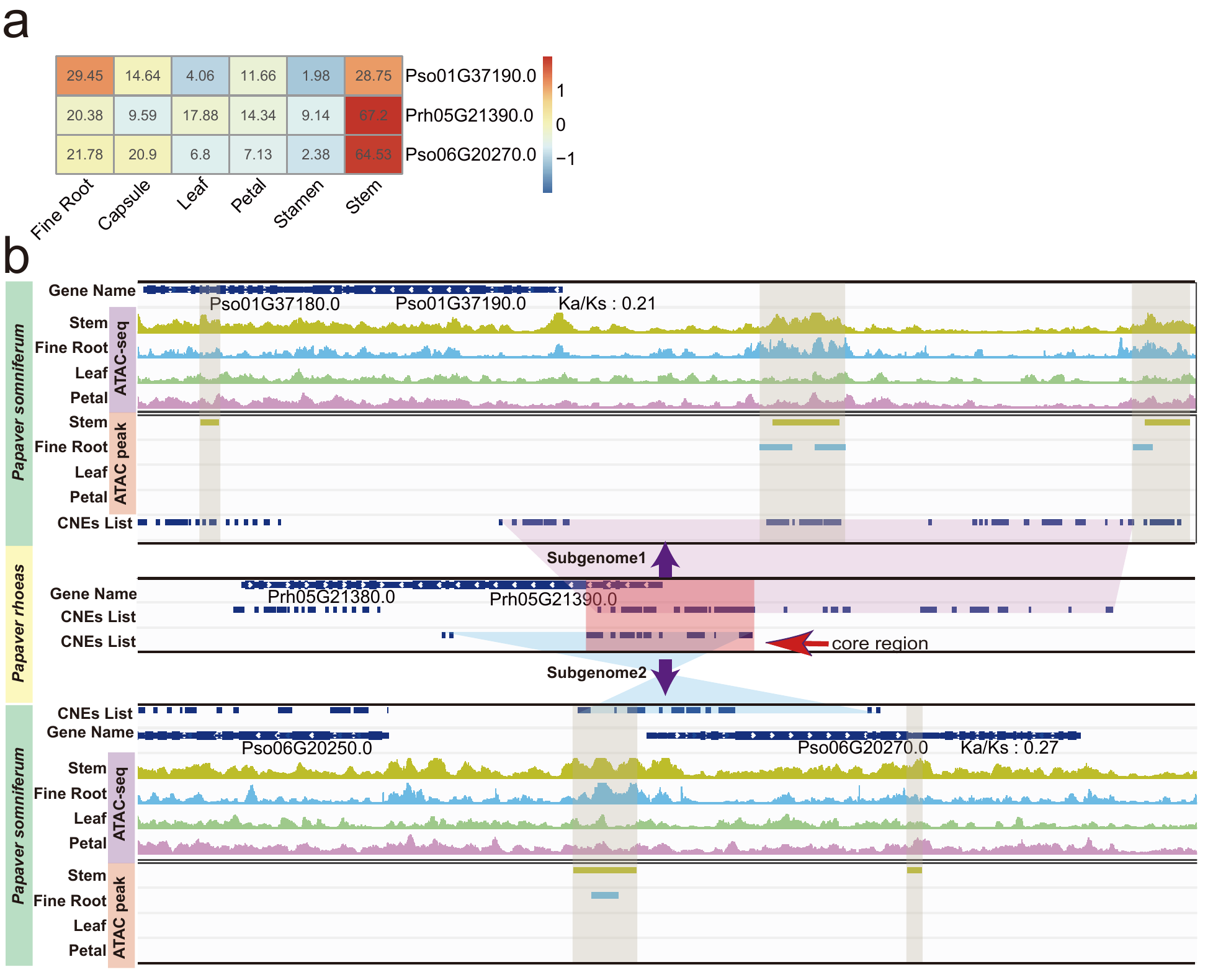

### Figure S16.tiff

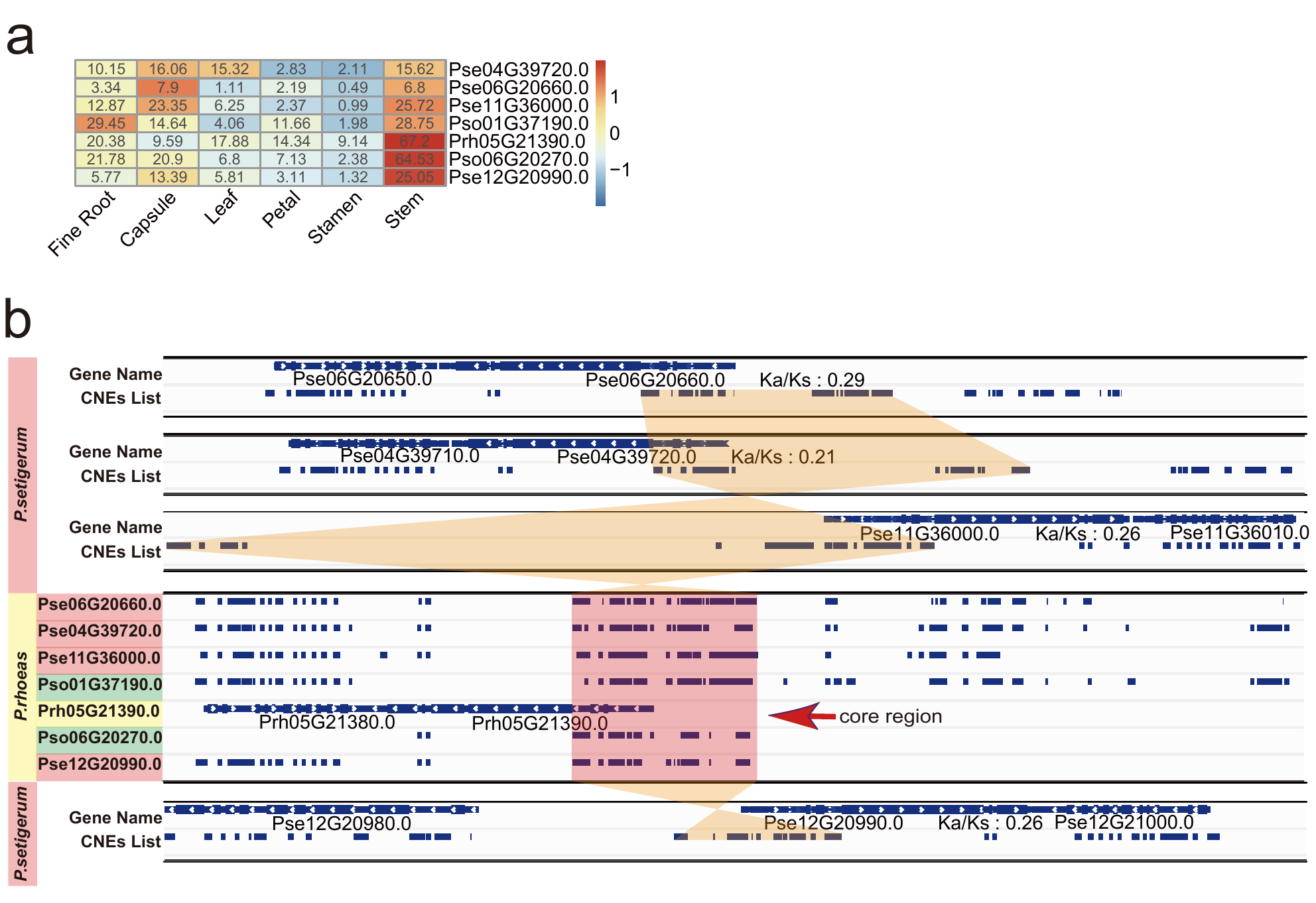

### Figure S17.tiff

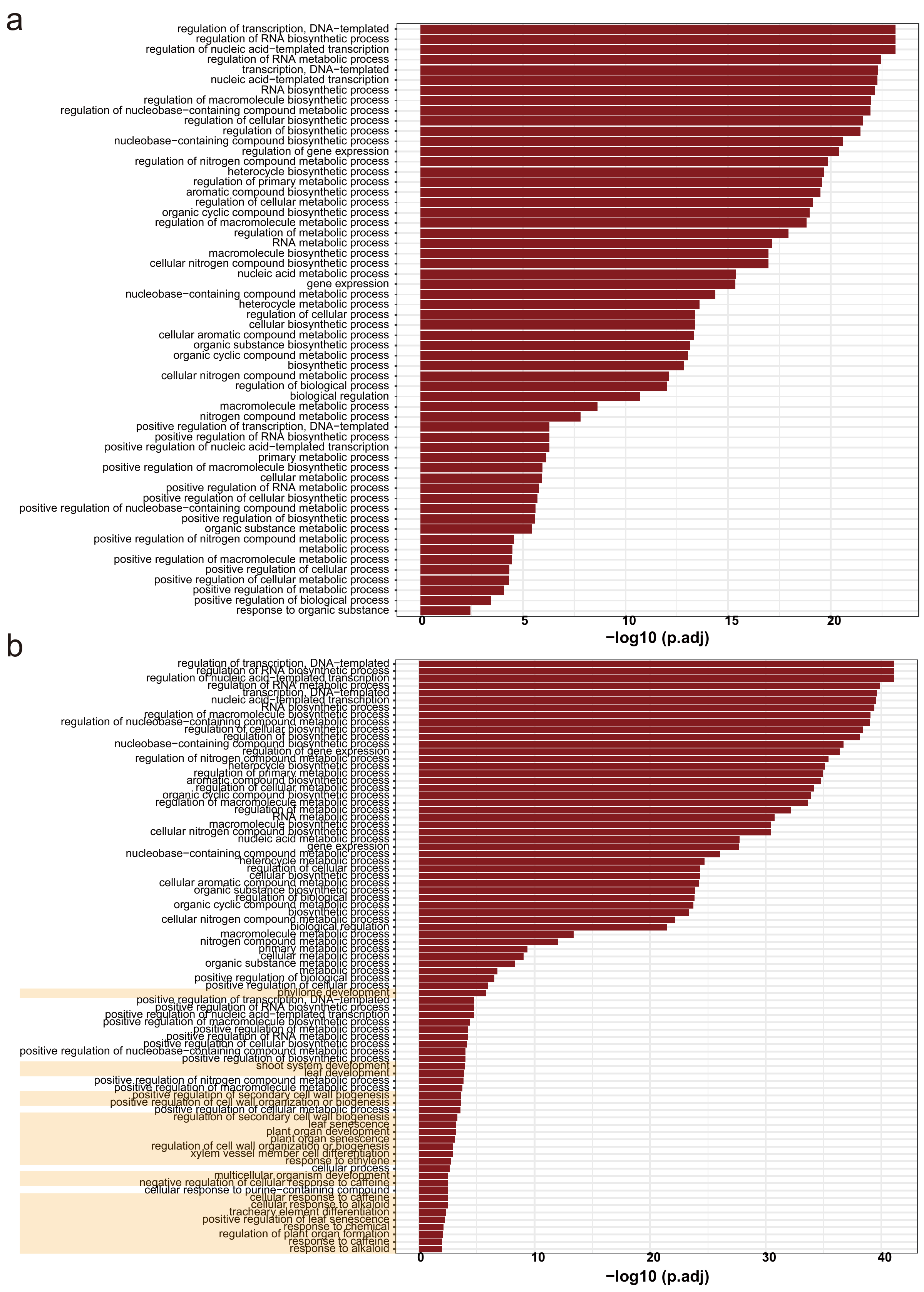

### Figure S18.tiff

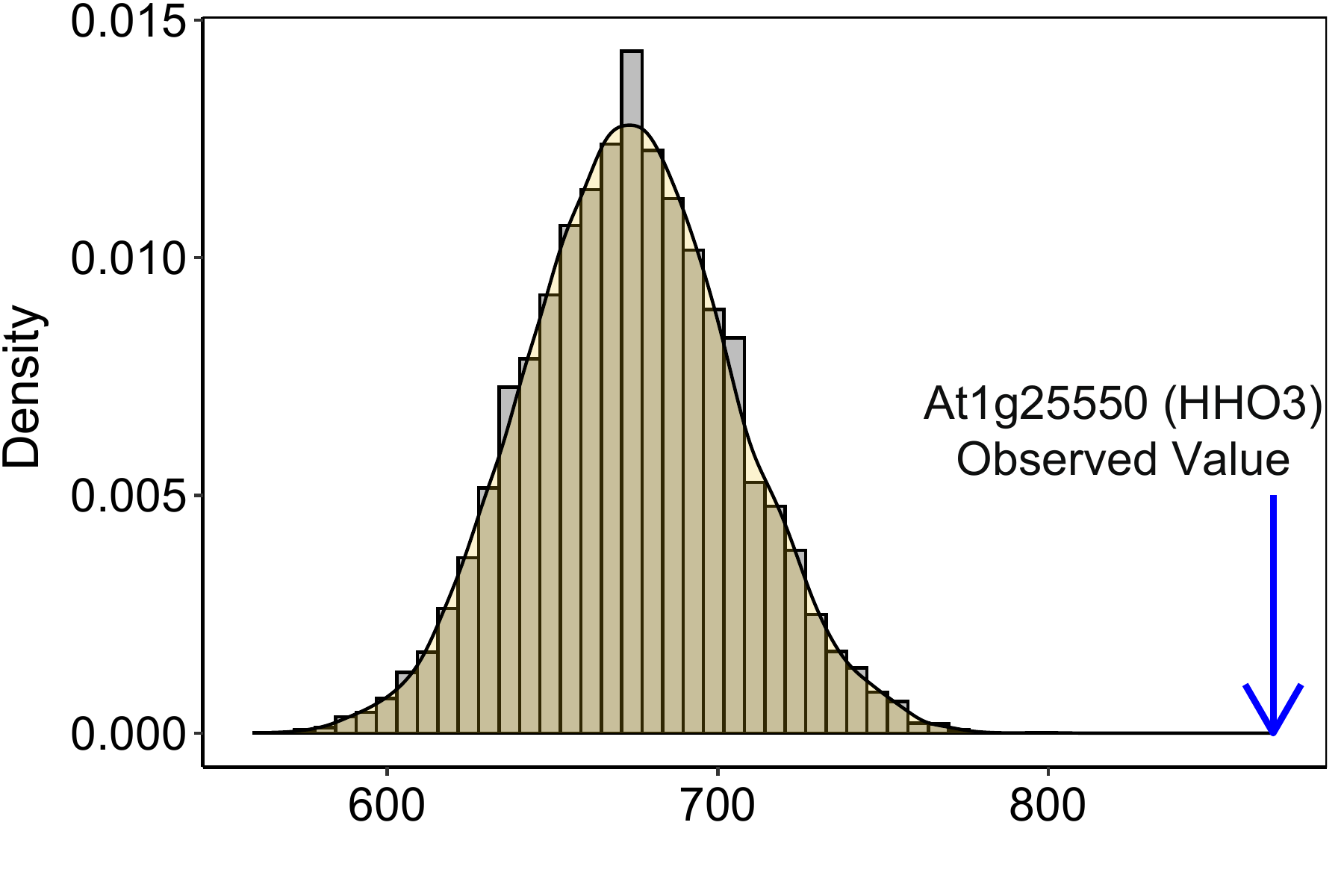

### Figure S19.tiff

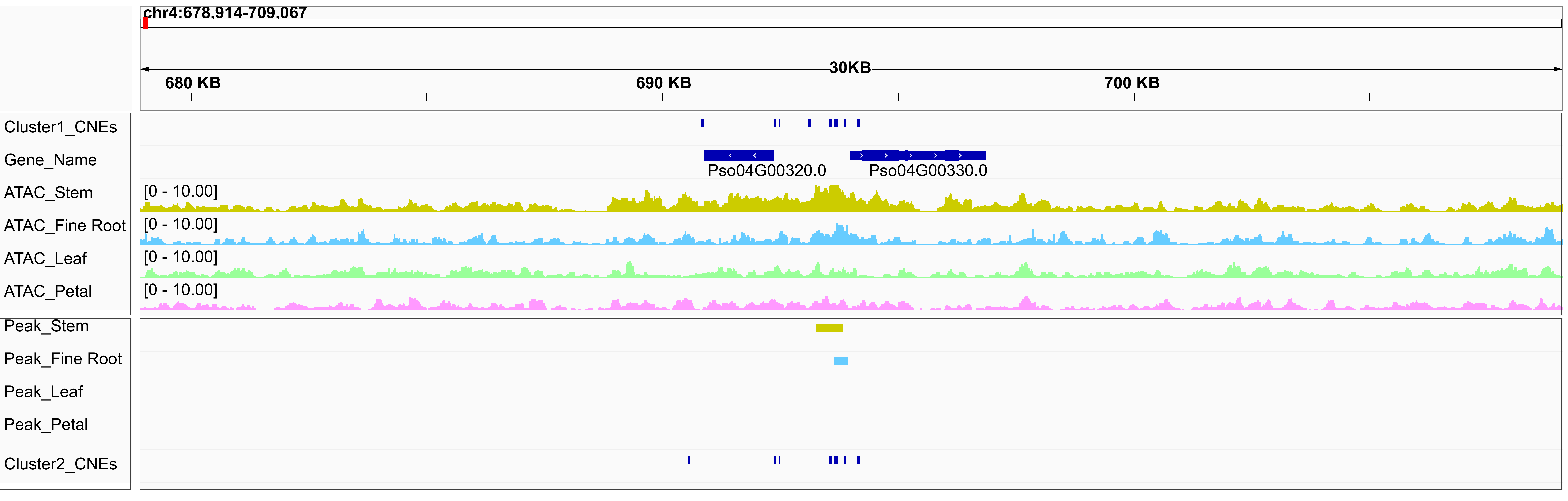
